# Structural mapping of antibody landscapes to human betacoronavirus spike proteins

**DOI:** 10.1101/2021.09.30.462459

**Authors:** Sandhya Bangaru, Aleksandar Antanasijevic, Nurgun Kose, Leigh M. Sewall, Abigail M. Jackson, Naveenchandra Suryadevara, Xiaoyan Zhan, Jonathan L. Torres, Jeffrey Copps, Alba Torrents de la Peña, James E. Crowe, Andrew B. Ward

## Abstract

Preexisting immunity against seasonal coronaviruses (CoV) represents an important variable in predicting antibody responses and disease severity to Severe Acute Respiratory Syndrome CoV-2 (SARS-2) infections. We used electron microscopy based polyclonal epitope mapping (EMPEM) to characterize the antibody specificities against β-CoV spike proteins in sera from healthy donors (HDs) or SARS-2 convalescent donors (CDs). We observed that most HDs possessed antibodies specific to seasonal human CoVs (HCoVs) OC43 and HKU1 spike proteins while the CDs showed reactivity across all human β-CoVs. Detailed molecular mapping of spike-antibody complexes revealed epitopes that were differentially targeted by antibodies in preexisting and convalescent serum. Our studies provide an antigenic landscape to β-HCoV spikes in the general population serving as a basis for cross-reactive epitope analyses in SARS-2 -infected individuals.

**One-Sentence summary:** We present the epitope mapping of polyclonal antibodies against beta-coronavirus spike proteins in human sera.

---

Four human coronaviruses (HCoVs) of genus α (HCoV-229E and HCoV-NL63) or β (HCoV-OC43 and HCoV-HKU1) are endemic in the human population contributing up to a third of the common cold infections (*1, 2*). While the infection rate and prevalence of these HCoVs vary based on the region, primary infections occur early in life with a majority of the population infected before 15 years of age (*2-5*). Most individuals possess antibodies to HCoVs targeting the trimeric spike glycoprotein and the nucleocapsid protein (N) though antibodies wane over time permitting reinfection even within a year (*6-9*). In addition to HCoVs OC43 and HKU1, the β-CoV genus also contains three highly pathogenic CoVs associated with human disease: Middle East Respiratory Syndrome CoV (MERS), Severe Acute Respiratory Syndrome CoV (SARS) and the novel SARS-2, the causative agent of the ongoing coronavirus disease 2019 (COVID-19) pandemic (*10, 11*).

The spike protein is an important determinant of host range and cell tropism as it mediates virus attachment and entry into the host cells, making it a major target for neutralizing antibodies and a key component for vaccine development (*12-16*). While the SARS-2 spike shares high structure and sequence homology with the SARS (69.2%) spike, it is less conserved across other β-CoVs, with as little as 27.2% sequence homology between SARS-2 and OC43 (*17*). Despite the low sequence conservation, preexisting immunity against seasonal CoV spike proteins has been associated with COVID-19 disease outcome as a consequence of either back-boost or induction of cross-reactive antibodies following SARS-2 infection (*8, 9, 18-22*). Of interest, COVID-19 donors with high SARS-2 antibody titers also possessed increased levels of antibodies against β-HCoVs (*8, 23*). It is not clear if infection triggers a recall of preexisting HCoV-specific antibodies or preferentially elicits cross-reactive β-CoV antibodies targeting conserved epitopes. Here, we elucidate the β-HCoV spike epitopes targeted by pre-existing serum antibodies and compare them to those elicited following SARS-2 infection using EMPEM methodology (*24, 25*)

Soluble ectodomains of spike proteins for β-CoVs, HKU1, OC43, SARS, MERS and SARS-2 (4 stabilized constructs were used for SARS-2) were generated and characterized by negative stain electron microscopy (ns-EM) and shown to be homogeneous in their prefusion conformation (Fig. S1A). To determine the baseline serum antibody titers to β-CoV spikes in the general population, either sera or plasma (based on availability) from eight HDs, collected prior to the COVID-19 pandemic, with unknown HCoV infection history were screened for spike antibodies by enzyme-linked immunosorbent assay (ELISA). All 8 donors exhibited reactivity to the OC43 spike, with half maximal effective concentration (EC_50_) serum dilution values ranging from 0.0007 to 0.02, while HKU1 antibody titers were lower in general (serum dilution EC_50_ of 0.001 to 0.06) (Fig. 1A and Fig. S1B). This finding is consistent with OC43 being the most commonly encountered HCoV globally while HKU1 is less prevalent (*4, 5*). Reactivity against SARS-2 spike was not detected in any of the HD sera, and only 1 out of 8 donors (D1124) exhibited low-level reactivity against SARS and MERS spikes. For comparison, we then assessed β-CoV spike reactivity in 3 sera samples from COVID-19 CDs (∼day 56 post-infection), all of whom exhibited high antibody titers to SARS-2 spike. Notably, the 3 donors also showed reactivity against other β-CoV spikes including SARS and MERS (Fig. 1A and fig. S1B). Given that the donors were immunologically naïve to these pathogenic CoVs, the data indicate that SARS-2 infection can elicit some level of cross-reactivity against the β-CoV spikes. While OC43 reactivity was high in both HD and CD sera, HKU1 spike antibody titers appeared enhanced in CDs (Fig. 1A and Fig. S1B). To investigate if serum antibody reactivity translated to inhibitory activity, we performed neutralization assays with both HD and CD serum against the OC43 virus and VSV-pseudotyped SARS and SARS-2 viruses. Overall, the serum inhibitory titers against the OC43 virus correlated well with their binding titers (Fig. 1A and Fig. S1C). While none of the HD sera neutralized SARS or SARS-2 virus, the CD sera exhibited neutralizing activity (Serum dilution IC_50_ of 0.002 to 0.007) against the SARS-2 virus and some weak activity against the SARS virus (Fig. 1A and Fig. S1C).

**Figure 1.**
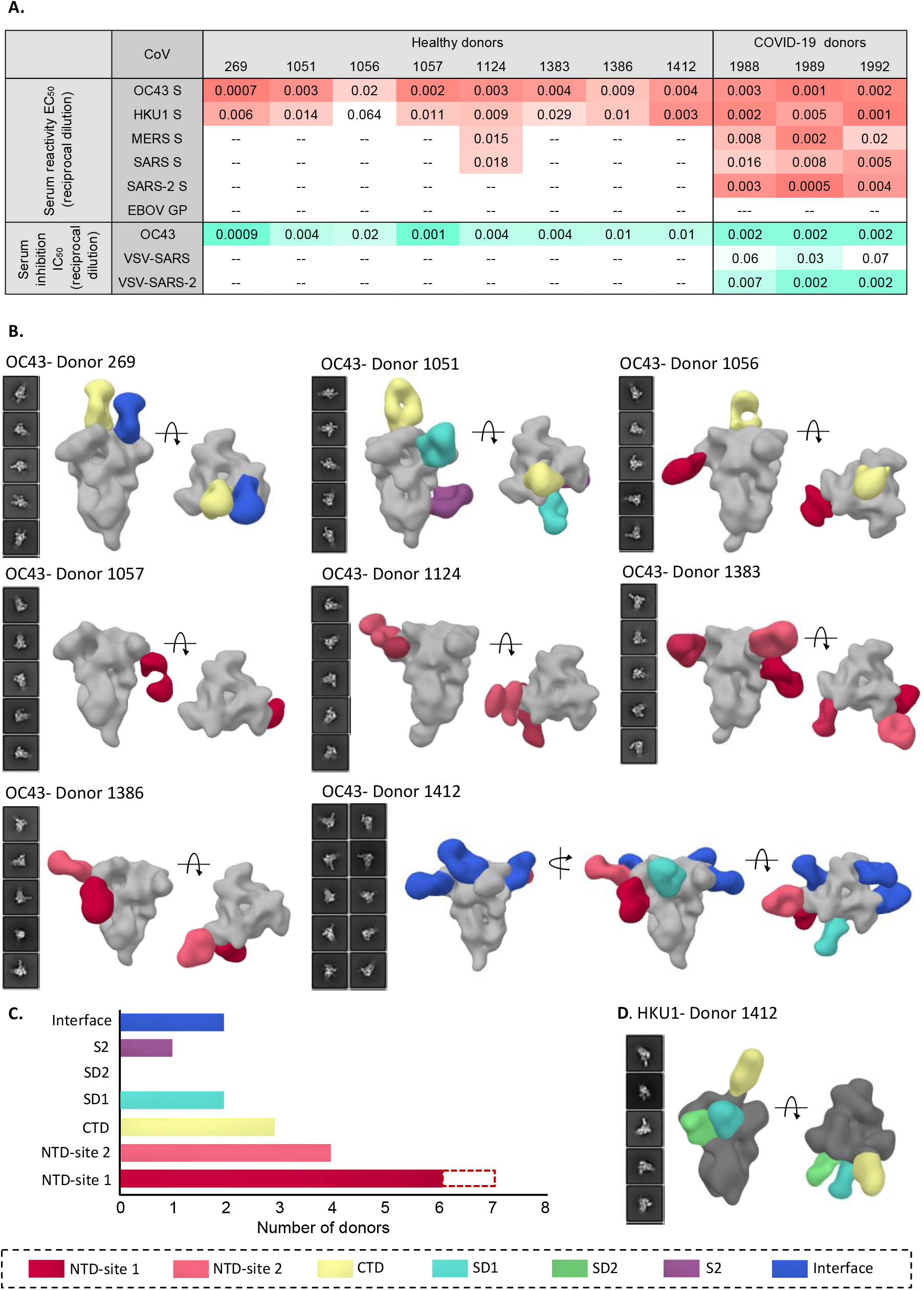
Human serum reactivity to β-CoV spikes. **(A)** ELISA half-maximal effective concentration (EC_50_) binding titers to OC43, HKU1, MERS, SARS and SARS-2 spikes and half-maximal inhibitory concentration (IC_50_) neutralization titers against OC43 virus and VSV-pseudotyped SARS or SARS-2 virus for HD and SARS-2 CD sera. Ebola virus glycoprotein (EBOV GP) was used as a negative control for detecting non-specific serum binding. Serum EC_50_ or IC_50_ titers are color-coded in gradients of orange or aquamarine, respectively. **(B)** Representative 2D classes and side and top views of composite figures from ns-EMPEM analysis of polyclonal Fabs from 8 HDs with the OC43 spike. **(C)** Bar graph summary of OC43 spike epitopes targeted by HD sera. Antibodies to NTD-site 1 were observed in 2D class averages for donor 269 but did not reconstruct in 3D, as indicated by dotted lines. **(D)** Composite figures from ns-EMPEM analysis of polyclonal Fabs from donor 1412 with the HKU1 spike. The Fab in panels and **(D)** are color-coded based on their epitope specificities as indicated at the bottom. OC43 or HKU1 spikes in panels **(B)** and **(D)** are represented in light gray or dark gray, respectively.

We next employed ns-EMPEM to determine the epitope specificities of spike antibodies in the HD sera. Structural analysis of polyclonal Fabs complexed with spike proteins from OC43, HKU1, SARS or MERS revealed OC43-reactive antibodies in all 8 donors and HKU1-reactive antibodies in 1 donor. We did not detect antibodies to either SARS or MERS spikes (Figs. 1B-D). Published cryo-EM structures of β-CoV spikes show the cleavable S1 and S2 subunits comprising an N-terminal domain (NTD), a C-terminal domain (CTD), subdomains 1 and 2 (SD1 and SD2), the fusion peptide (FP) and heptad repeats 1 and 2 (HR1 and HR2) (*26-29*). OC43 NTD-reactive Fabs were seen in 7 donors targeting either the 9-O-acetylated sialic acid receptor binding site (RBS) defined by loop residues 27 to 32, 80 to 86, 90 and 95 (NTD-site 1) or a site adjacent to the RBS encompassing residues from loops 112 to119, 176 to 186 and 254 to 261 (NTD-site 2) (Figs. 1B-C). The prevalence of NTD-site 1 Fabs that can sterically block receptor engagement correlated well with the OC43 inhibitory titers observed across HDs. While neither the CTD nor the SD1 of OC43 spike is associated with any known function, antibodies to CTD were seen in at least 3 donors, to SD1 in 2 donors and to S1 inter-protomeric interfaces in 2 donors. A single S2-reactive antibody from donor 1051 displayed a broad footprint with potential interactions with residues 800 to 807, 1013 to 1031, and 1062 to 1068 (Fig. S1D). Of interest, donor 1412 with a relatively low OC43 neutralization titer displayed the greatest diversity of Fab specificities, targeting 6 distinct S1 epitopes including the inter-protomeric interfaces. This individual was also the only donor with detectable Fab responses to the HKU1 spike targeting the CTD, the SD1 and the SD2 (Fig. 1D).

Samples from 3 donors (269, 1051 and 1412) were chosen for high resolution cryo-EMPEM studies with OC43 spike as they represented individuals with antibodies against all the unique epitopes observed (Table S1). We reconstructed 10 high-resolution maps of unique spike-Fab complexes (Fig. 2A, Figs. S2-5, Table S2). High-resolution analysis of immune complexes from donor 269 revealed Fabs bound to the CTD, CTD-NTD interface and to NTD-site 1; NTD-site 1 Fab was not reconstructed during the ns-EMPEM studies. For donor 1051, cryo-EM analysis enabled differentiation of polyclonal Fabs targeting the CTD that were originally observed as a single species by ns-EM. We were unable to obtain reconstructions of either the SD1 or S2 antibody despite multiple attempts at focused classification, likely owing to low Fab abundance or dissociation of the complex during the cryogenic sample preparation process. For donor 1412, we reconstructed 5 of the 6 specificities seen in ns-EMPEM, targeting NTD-site 1, NTD-site 2, SD and inter-protomeric S1 interfaces. In all reconstructed maps we observed an additional non-spike density buried within a hydrophobic pocket in the CTD; the location and size resembling linoleic acid in SARS-2 spike (*30, 31*). The MW of 254 g/mol obtained by mass-spectrometry analysis of the OC43 spike and the corresponding density in the OC43 map are however consistent with sapienic acid (6Z-hexadecenoic acid; Fig. 2B and Fig. S5A). The aliphatic chain of sapienic acid improves the hydrophobic packing at the CTD-CTD interface of two adjacent protomers, while the carboxyl group forms hydrogen bonds with the side chain of Tyr395 in the CTD of one protomer and the main chains of residues Leu422 and/or Gly423 within the CTD of the other protomer (Fig. 2B). These contacts are likely helping stabilize the closed conformation of OC43 spike.

**Figure 2.**
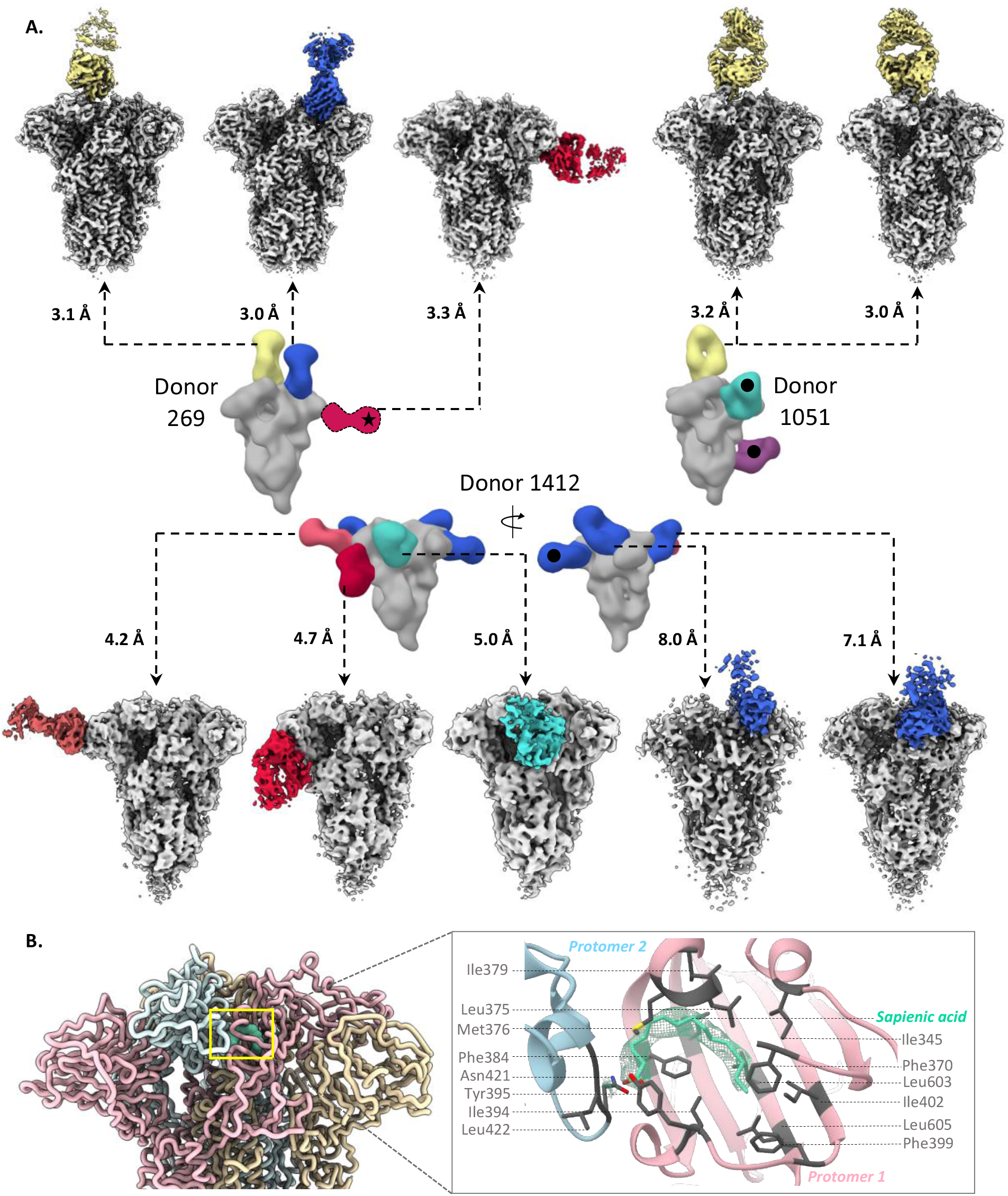
CryoEMPEM analysis of OC43 spike-polyclonal Fab complexes. **(A)** High-resolution cryo-EMPEM reconstructions of OC43 spike complexed with polyclonal Fabs derived from sera from HDs 269 (top-left), 1051 (top-right) or 1412 (bottom); the representative composite figures from ns-EMPEM from these donors are shown in the middle. Each map depicts a structurally unique polyclonal antibody class reconstructed at the indicated resolution with the Fabs colored according to the scheme used in Fig. 1. OC43 spike is represented in light gray. Fabs marked with a black dot were observed by ns-EMPEM but were not detected by cryo-EMPEM. Fab class from donor 269 marked with a star was resolved by cryo-EMPEM but not by ns-EMPEM. **(B)** Sapienic acid (aquamarine) binding within a hydrophobic pocket in the CTD-CTD inter-protomeric interface. Protomers are colored in light pink, blue or wheat and the interacting residues are shown in gray.

Atomic models of spike-Fab complexes were relaxed in 7 out of 10 maps with resolutions ≤ 4.8Å (Fig. 3A-G, Figs. S6-7). Fabs were represented as poly-alanine pseudo-models. Both Fabs to immunodominant NTD-site 1 (Fab1-spike at 3.3 Å and Fab2-spike at 4.7 Å), approach the RBS at different angles with dissimilar engagement of antibody heavy and light chain complementarity determining region (HCDR and LCDR; Figs. 3A-B and Figs. S6A-B). While Fab1 made spike contacts at residues 33 to 36 (using HCDR2), 39 to 42 (HCDR3), 88 to 89 (LCDR3) and 264 to 267 (LCDR1), Fab2 approaches at a much steeper angle by inserting its HCDR3 into the NTD pocket encompassing loops 82 to 86, 35 to 43, and 263 to 270 along with other LCDR1 and LCDR2 contacts at residues 40 to 44. Fab3 (4.2 Å), targeting the second prevalent site, NTD-site 2, binds adjacent to RBS with main contacts at residues 118 to 121 (LCDR3) and interacting with loops 183 to 187 (HCDR3) and 261 to 265 (LCDR1) (Fig. 3C and Fig. S6C). While antibodies to NTD-site 1 directly overlap with the RBS, NTD-site 2 antibodies could potentially block receptor binding by steric hindrance. Collectively, cryo-EMPEM analysis of Fabs to NTD reveal structural features of these immunodominant epitopes associated with anti-viral activity against OC43.

**Figure 3.**
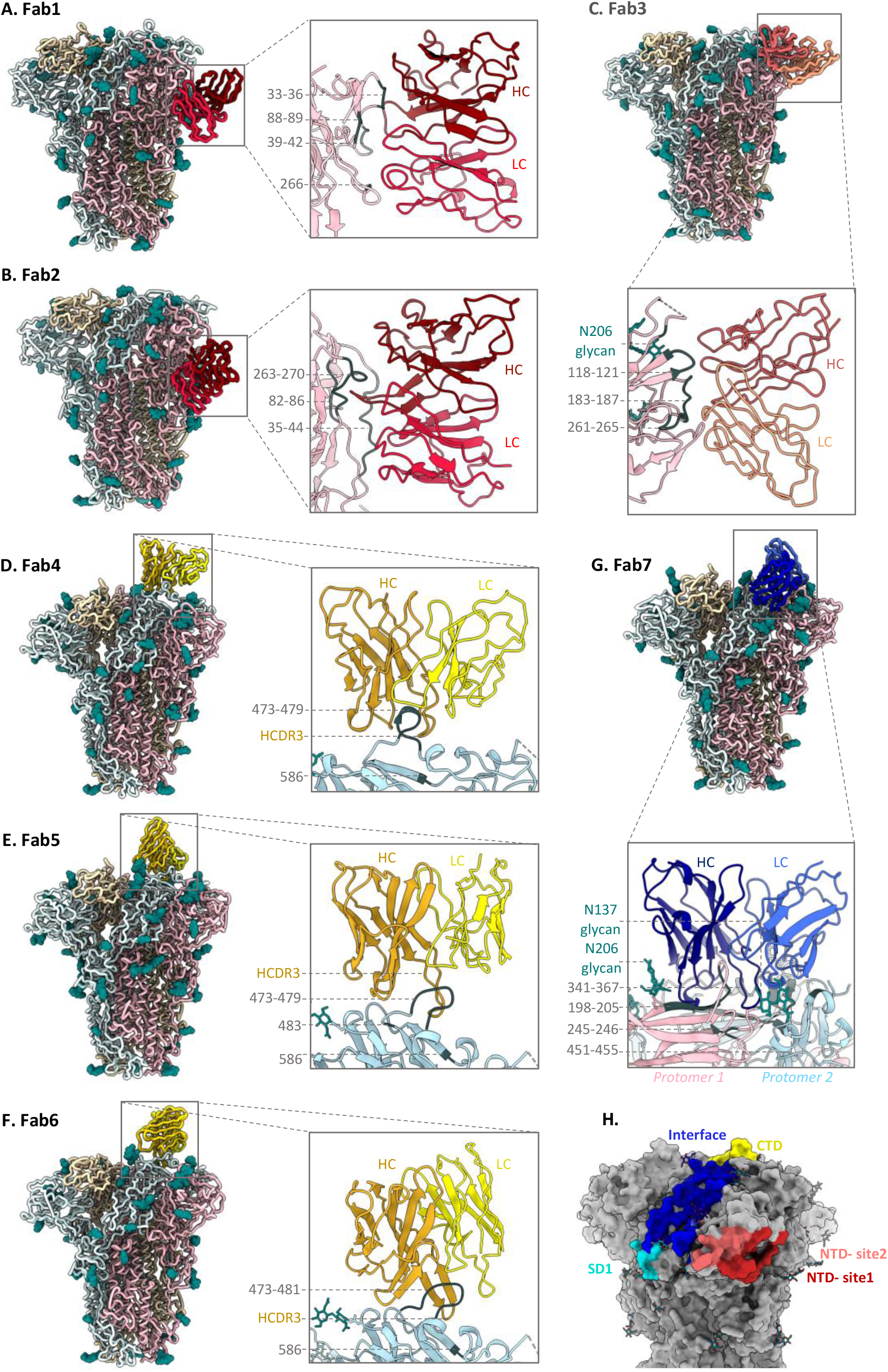
Cryo-EM structures of polyclonal Fabs targeting the OC43 spike. **(A-G)** Tube or ribbon representation of atomic models of OC43 spike-Fab complexes along with zoomed-in views of epitope-paratope interactions. **(A-B)** Fab1 and Fab2 (red) target the NTD-site1 or RBS, Fab3 (orange) targets NTD-site2 adjacent to RBS, **(D-F)** Fab4, Fab5 and Fab6 (yellow) target the CTD and **(G)** Fab7 (blue) targets the NTD-CTD interface. The spike protomers are shown in light blue, light pink or wheat (ribbon representation) with glycans in teal (sphere atom representation) and primary epitope contacts in gray. Detailed contact residues along with corresponding EM densities are shown in Figs. S6-7. **(H)** Surface representation of OC43 spike (gray) showing collective epitopes of Fab1 to Fab10 colored based on their binding site using the color scheme from Fig. 1.

High resolution reconstructions of 3 CTD Fab-spike complexes (Fab4 at 3.1 Å, Fab5 at 3.2 Å and Fab6 at 3.0 Å reveal an almost identical epitope featuring a single CTD loop 472 to 483 (Figs. 3D-F and Figs. S6D-F). Fab4 surrounds the loop with HCDR1, HCDR2, HCDR3 and LCDR3 making contacts with residues 473 to 479, and Trp586. Whereas Fab4 binding did not induce any conformational changes in the loop residues 472-483 in comparison to the published OC43 apo-spike structure (PDB# 6OHW, (*29*)), both Fab5 and Fab6 stabilize the loop in a different conformation (Figs. 3D-F and Figs. S6D-F). While Fab5 uses HCDR2, HCDR3, LCDR1 and LCDR3 to interact with loop residues 474-477 and 483 with potential HCDR3 contact at Trp586, Fab6 binds in a similar manner with the main distinguishing features being HCDR2 interaction with Thr481 instead of His483, additional HCDR1 contact with Val479 and the displacement of glycan at position Asn449 by the longer Fab6 LCDR1 (Fig. 3E-F and Figs. S6E-F). Overall, structural analysis of antibodies to CTD reveal the loop 472 to 483 as the major antigenic element that is generally sandwiched between multiple CDRs with Trp586 stabilizing the interaction.

Among the 3 interprotomeric antibodies reconstructed, Fab7 (3 Å) and Fab8 (8 Å) bind two structural domains (CTD and NTD) while Fab9 (7.1 Å) extends across 3 domains (NTD, CTD and SD1; Fig. 3G and Figs. S7A-C). Fab7 interacts extensively with both NTD (residues 140, 169, 198, 200 to 205, 245 to 246) and CTD (residues 341 to 367, 451 to 455 and 469) using all CDRs, although the main interactions are facilitated by HCDR3. The antibody makes glycan contacts at position Asn206 while shifting the Asn137 glycan from its original position to accommodate LCDR1 (Fig. 3G and Fig. S7A). Fab8 epitope, comprised of NTD loop 196 to 208 and CTD loop 339 to 352, is bordered by 4 glycans with potential contacts with glycans at positions Asn206 and Asn449 and the Fab9 interaction is driven by NTD loops 204 to 210 and 149 to 159 on the first protomer with potential Asn206 glycan contact and by two CTD loops 337 to 341 and 366 to 367 and two SD1 loops 672 to 676 and 622 to 624 on the adjacent protomer (Figs. S7B-C). Notably, the Asn675 glycan is buried in the Fab9 spike interface making contact with the antibody (Fig. S7C). Lastly, Fab10 (5 Å) targets the SD1 with primary interactions with Asp624, Glu646, Arg676, and glycans at Asn648 and Asn678 (Fig. S7D). Importantly, Fab9 and Fab10 both make extensive contacts to the Asn675 glycan which represents an important immunogenic determinant within the SD1 epitope. An epitope summary of commonly elicited β-CoV spike antibodies in healthy human serum is shown in Fig. 3H and CDR lengths for Fabs1-7 determined by structural homology are summarized in table S3

Next, we sought to investigate the nature of spike antibodies in serum following SARS-2 infection. Sera from 3 CDs were screened for antibodies to SARS-2 spike by ns-EMPEM. Analysis of EM data (2D and 3D) revealed both NTD and RBD (or CTD) antibodies though the latter were relatively fewer in number and more difficult to reconstruct owing to the flexible RBD (Fig. 4A). While RBD antibodies have been well documented to provide protection against SARS-2 infection, NTD responses are being recognized as an important component of the neutralizing response to SARS-2, particularly those targeting the supersite comprising of residues 14 to 20, 140 to 158 and 245 to 264 (*32-37*). Collectively, these donors possessed several polyclonal antibodies targeting this supersite along with antibodies to other previously described sites (*32*). Interestingly, we also observed antibody-pairs in both donors 1988 and 1989 that appeared to partially bind each other while also recognizing some part of the spike NTD (Fig. 4A). It is unclear why these antibodies are triggered in SARS-2 donors, and the implications of this finding for understanding COVID pathology needs further investigation.

**Figure 4.**
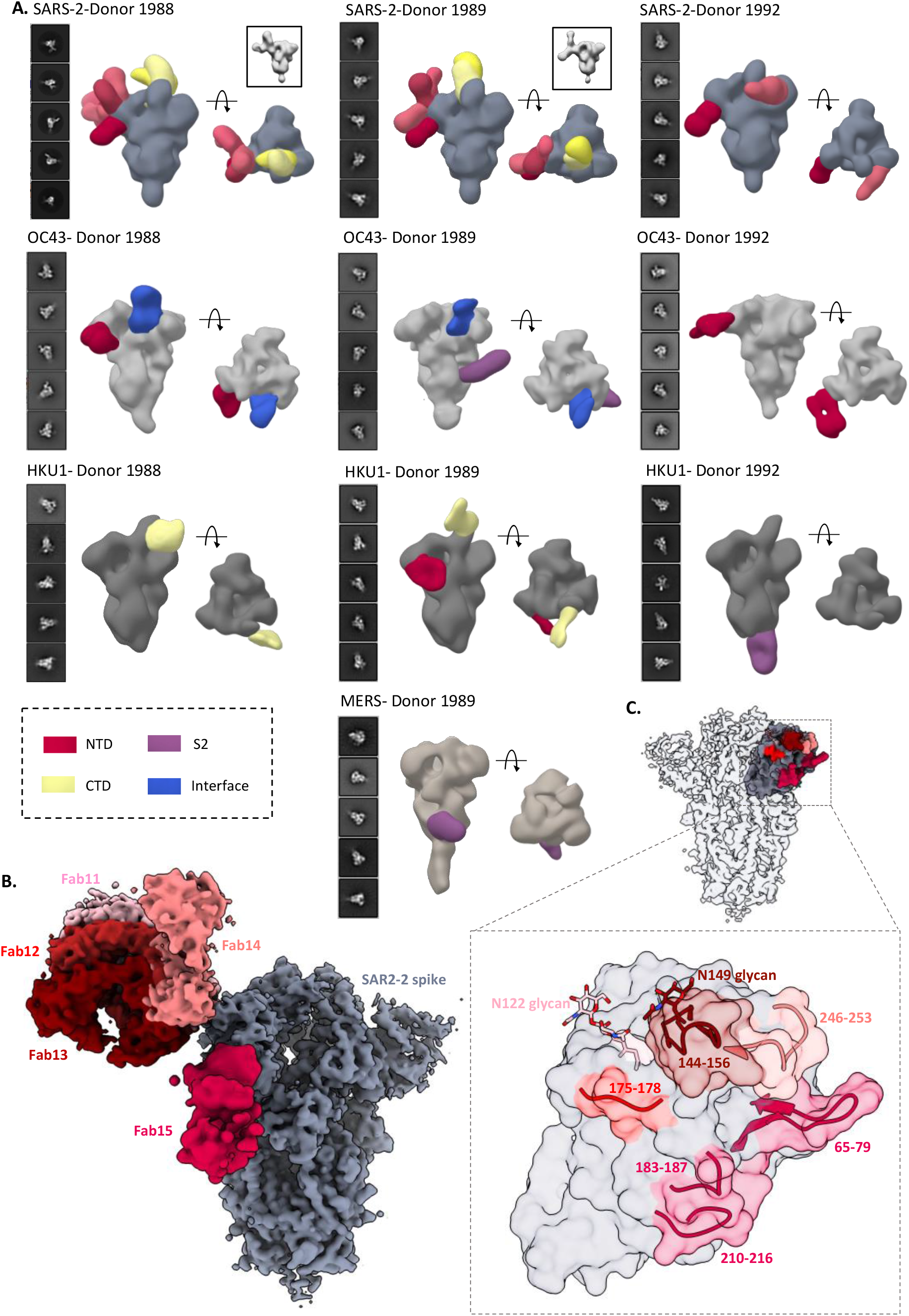
Ns- and cryo-EMPEM analysis of polyclonal Fabs from SARS-2 CD sera. **(A)** Representative 2D classes and side and top views of composite figures from ns-EMPEM analysis of polyclonal Fabs from 3 SARS-2 donors complexed with β-CoV spikes. The donor numbers along with the corresponding CoV spikes are indicated above each panel in **(A)**. The Fabs are color-coded based on their epitope specificities as indicated at the bottom-left. SARS-2, OC43, HKU1 and MERS spikes are represented in slate gray, light gray, dark gray and beige respectively. 3D reconstructions displaying potential self-reactive antibodies are shown in grey on the top right corners for both donors 1988 and donor 1999 in complex with SARS-2 spike **(B)** Composite figure showing 5 unique antibody classes, Fab11 to Fab15 colored in shades of red, to SARS-2 spike NTD reconstructed using cryo-EMPEM analysis of polyclonal Fabs from donors 1988 and 1989 complexed with SARS-2 stabilized spikes. **(C)** Surface representation of SARS-2 spike showing epitopes of Fabs 11 to 15 from **(B)** on a single NTD (slate gray) with a zoomed-in view displaying the loop residues comprising each epitope. Loop 144 to 156 with the N149 glycan forms an immunodominant element commonly targeted by Fabs 11 to 14. The sub-epitope colors correspond to each Fab shown in **(B)**

To obtain detailed molecular information on immunodominant epitopes within the SARS-2 spike, we subjected polyclonal samples from 2 CDs (pooled Fabs from donors 1988 and 1989) to cryo-EMPEM analysis with SARS-2 spike. The analysis yielded 5 maps featuring NTD antibodies (Fab11 to Fab15) (Figs. 4B-C, Figs. S8-9 and Table S4). Fabs 11 to 14, were reconstructed at resolutions 3.9 Å, 4.2 Å, 4.3 Å or 4.4 Å, respectively, and were all immunofocused onto the NTD loop 145 to 155 with the Asn149 glycan present at the core of each interaction (Figs. 4B-C). Fab11 with its tilted angle of approach also made contacts with the Asn122 glycan while Fab12 and Fab13 appeared to make some additional contacts with adjacent loops 176 to 181 and 247 to 252. Although Fab14 also binds to loop 145 to 155, its distinctly different angle of approach also allows extensive contacts with loop 246 to 253, similar to supersite antibodies (*32*). In contrast, Fab15 interacts with loops 65 to 79, 183 to 187 and 210 to 217, an antigenic site similar to that recognized by the antibody S2M24 (*32*).

Our ELISA data demonstrated that there is an increase in antibody binding titers to non-SARS-2 β-CoV spikes following an infection with SARS-2 virus (“convalescent donors”) (Fig 1A). This finding indicates either a back-boost of pre-existing responses or elicitation of novel cross-reactive antibodies to conserved epitopes. Structural mapping of SARS-2 spike residues that are either identical to or have a conserved substitution in at least 3 of the 4 other β-CoVs, OC43, HKU1, SARS and MERS, revealed several conserved patches in the S2 subunit that could potentially elicit cross-reactive responses (Fig. S10A). Several recent studies have found S2 as a target for cross-reactivity across β-CoVs (*8, 9, 18, 19, 22*). Two individual studies also revealed SARS-2 spike residues in and around 560 to 572, 819 to 824, and 1,150 to 1,156 and their homologous regions on other HCoV spikes as being recognized with higher frequency in COVID-19 patients as compared to pre-COVID controls (Fig. S10A)(*18, 20*). To determine if these epitopes are targeted following SARS-2 infection, we performed ns-EMPEM on CD sera with OC43, HKU1, SARS and MERS. As with HDs, the SARS-2 CDs had serum antibodies to the OC43 spike protein; antibodies to NTD-site 1 were seen in 2 donors, antibodies to interface in 2 donors and an S2 antibody was observed in 1 donor (Donor 1989) (Fig. 4A). While we are uncertain if the S2 antibody was induced by SARS-2 infection, the antibody appears to target the helix 1014 to 1030 that is highly conserved across the β-CoV spikes (Fig. S10B). Notably, donors who possessed high levels of OC43 antibodies also had some SARS-2-reactive antibodies pre-pandemic that did not correlate with protection against SARS-2 (*9*). When complexed with the HKU1 spike, we were able to detect antibodies in all 3 CD samples, which was higher than seen for HD (3D reconstructions were possible only for 1 of the 8 HD sera) suggesting an increase in HKU1 antibody titers following SARS-2 infection (Fig. 4A). Of interest, Song *et al*. observed higher HKU1 spike antibody titers in post-COVID sera compared to pre-pandemic sera, whereas titers remained comparable for other HCoV spikes (*8*). Whereas donors 1988 and 1989 had antibodies to the HKU1 CTD and/or the NTD, donor 1992 sera contained an S2 antibody binding to the base of the trimer. The epitope is analogous to that of the β-CoV cross-reactive spike monoclonal antibody (mAb) CC40.8 isolated from a COVID donor (Fig. S10C); CC40.8 binds strongly to SARS-2 and HKU1 spikes while also exhibiting some reactivity to SARS and OC43 spikes (*8*). We also reconstructed a MERS spike antibody in donor 1989 that partly overlaps with the known MERS mAb G4 targeting the S2 connector domain near the trimer base (Fig. S10D)(*27*). The presence of a MERS-reactive antibody in a MERS-naive donor illustrates induction of cross-reactive responses following SARS-2 infection. We were not able to reconstruct any antibodies to SARS even though the CD sera had detectable titers against the spike. An overall comparison of antibody specificities between the HD and CD sera revealed antibody classes that were present in both the groups primarily targeting the S1 subunit while antibodies to the more conserved S2 subunit were enriched in the COVID donors. Collectively, these results suggest that SARS-2 infection triggers induction of cross-reactive antibodies to conserved β-CoV spike epitopes while some HCoV spike-specific antibodies may be back-boosted. This cross-boosting while associated in COVID-19 pathogenesis may also have long-lasting implications for immunity to seasonal CoVs as much of the population will be vaccinated and/or infected with SARS-2.

## Acknowledgements

We thank Bill Anderson, Hannah L. Turner and Charles A. Bowman for their help with electron microscopy, data acquisition and data processing. We thank Bill Webb and Linh Truc Hoang for their assistance with mass spectrometry and data processing. We thank Lauren Holden for her assistance with the manuscript.

## Author contributions

SB, ABW and JEC conceived and designed the study. SB, AA, NK, LMS, AMJ, NS, XZ, JLT, JC and ATdlP performed experiments. SB and AA performed cryo-EMPEM, model building and refinements. SB, AA, NK, ABW and JEC provided intellectual contributions. SB and ABW wrote the paper and all authors reviewed and approved the final version of the manuscript.

## Data and materials availability

Cryo-EM maps have been deposited at the Electron Microscopy Data Bank (EMDB) with accession codes EMD-24968 (OC43 spike-Fab1), EMD-24969 (OC43 spike-Fab2), EMD-24970 (OC43 spike-Fab3), EMD-24989 (OC43 spike-Fab4), EMD-24990 (OC43 spike-Fab5), EMD-24991 (OC43 spike-Fab6), EMD-24992 (OC43 spike-Fab7), EMD-24993 (OC43 spike-Fab8), EMD-24994 (OC43 spike-Fab9), EMD-24995 (OC43 spike-Fab10), EMD-24996 (SARS-2 spike-Fab11), EMD-24997 (SARS-2 spike-Fab12), EMD-24998 (SARS-2 spike-Fab13), EMD-24999 (SARS-2 spike-Fab14), EMD-25000 (SARS-2 spike-Fab15). Atomic models have been deposited to the RCSB protein data bank (PDB) with PDB IDs: 7SB3 (OC43 spike-Fab1), 7SB4 (OC43 spike-Fab2), 7SB5 (OC43 spike-Fab3), 7SBV (OC43 spike-Fab4), 7SBW (OC43 spike-Fab5), 7SBX (OC43 spike-Fab6), 7SBY (OC43 spike-Fab7). The EMDB accession codes for ns-EM apo spikes HKU1, OC43, MERS, SARS, SARS-2 HP-GSAS, SARS-2 HP-GSAS mut2, SARS-2 HP-GSAS mut4 and SARS-2 HP-GSAS mut7 are EMD-25014 – EMD-25021 respectively. The EMDB accession codes for ns-EMPEM polyclonal datasets are EMD-25001 (Donor 1051), EMD-25002 (Donor 1056), EMD-25003 (Donor 1057), EMD-25004 (Donor 1124), EMD-25005 (Donor 1383), EMD-25006 (Donor 1386), EMD-25009 (Donor 1412), EMD-25010 (Donor 269), EMD-25011 (Donor 1988), EMD-25012 (Donor 1989) and EMD-25013 (Donor 1992)

## SUPPLEMENTARY MATERIALS

Materials and Methods

Figures S1 to S10

Table S1 to S4

References (1-25)

