## Supplemental Material for "Structural mapping of antibody landscapes to human betacoronavirus spike proteins"


**This PDF file includes:**

Materials and Methods
Figures S1 to S10
Tables S1 to S4
References

#### Materials and methods

##### Expression and purification of recombinant spike proteins

All spike ectodomain constructs contain a C-terminal T4 fibritin trimerization domain, an HRV3C cleavage site, an 8xHis-Tag and a Twin-strep-tag for purification. The HKU1 spike construct includes the residues 1 to 1,276 from isolate N5 (GenBank Q0ZME7) with the S1/S2 cleavage site modified to 752-GGSGS-756 and the residues 1,067 to 1,068 replaced by prolines for generating stable uncleaved spike proteins. The OC43 spike construct contains spike residues 1 to 1,287 (GenBank AIL49484.1) with introduction of stabilizing prolines at sites 1,079 and 1,080. The SARS spike construct was generated with residues 1to 1,196 of the Tor2 strain (GenBank AAP41037.1) with stabilizing prolines at residues 968 and 969 while the MERS construct was synthesized with residues 1 to 1,291 from the England1 strain (GenBank AFY13307.1) with stabilizing prolines at positions 1,060 to 1,061 and the S1/S2 cleavage site modified to 748-ASVG-751. For the SARS-2 spike, we synthesized a base construct (HP-GSAS) with residues 1 to 1,208 from the Wuhan-Hu-1 strain (GenBank: QHD43416.1) with 6 stabilizing proline (HexaPro) substitutions at positions 817, 892, 899, 942, 986 and 987 and the S1/S2 furin cleavage site modified to 682-GSAS-685. We also generated 3 other HP-GSAS constructs each with a pair of cysteine substitutions to generate stable disulphide linkages: HP-GSAS Mut2 (S383C and D985C), HP-GSAS Mut4 (A570C and L966C) and HP-GSAS Mut7 (V705C and T883C). We used an equal ratio mixture of all 4 spikes for each assay (*1, 2*).

For protein expression, FreeStyle 293-F cells were transected with the spike plasmid of interest and cultures were harvested at 6-days post-transfection. For OC43, HKU1, SARS and MERS, the spike proteins were purified from the supernatants on cOmplete™ His-Tag Purification Resin (Millipore Sigma) using a 250 mM imidazole elution buffer and buffer exchanged to Tris-NaCl buffer (25 mM Tris, 500 mM NaCl, pH 7.4) before further purification with Superose 6 increase (S6i) 10/300 column (GE Healthcare Biosciences). For SARS-2 spikes, we used StrepTactin-XT 4FLOW high capacity columns (IBA Lifesciences) and elution with buffer BXT (IBA Lifesciences) prior to buffer exchange with buffer W (IBA Lifesciences) and S6i column purification. Protein fractions corresponding to the trimeric spike proteins were collected and concentrated. The quality of purified proteins was assessed by negative stain EM.

**Human samples used in the study**

For all the assays described in the paper, serum samples were used for donors 269, 1051, 1056, 1057, 1124, 1383 and 1412 while plasma samples were used for donors 1988, 1989 and 1992. The terms serum and plasma are used interchangeably in the manuscript.

**ELISA for evaluating serum reactivity to spike proteins**

To determine the half-maximal effective concentration ($EC_{50}$) binding titers for donor sera, we performed ELISA using 384-well plates that were coated overnight with 1 µg/mL of recombinant spike protein of interest and subsequently blocked with 5 % non-fat dry milk and 2 % goat serum in PBST (PBS with 0.1 % Tween-20) for 1 hr at RT. Plates were washed and 25 µL of two-fold serially diluted sera starting with a four-fold dilution were added to the wells and incubated for 1 hr. The washed plates were incubated with anti-human IgG alkaline phosphatase conjugate (Meridian life science) for 1 hr and with 25 µL of phosphatase substrate solution following a final wash. The optical density values were measured at 405 nm wavelength following a 20 min incubation and the corresponding $EC_{50}$ values were calculated using Prism software (GraphPad) using non-linear regression analysis.

**HCoV-OC43 serum neutralization assay**

HCT-8 cells were seeded in 96 well plates at a density of 10,000 cells/well in Gibco RPMI-1640 medium with 10% FBS (ThermoFisher Scientific) and incubated overnight. Heat-inactivated serum was diluted in RPMI-1640 medium and incubated with OC43 virus for 1 hr prior. 50 µL of the mixture was added to each well of the 96-well plate containing the HCT-8 cells and incubated again for 1 hr prior to addition of 100 µL RPMI-1640 to each well. The plates were incubated for 4 days at 37°C and the supernatants were harvested to perform hemagglutination inhibition assay. 50 µL of supernatant was mixed with 50 µL of turkey red blood cells and plated on v-bottom plates. The hemagglutination results were recorded after 30 min of incubation.

**Real-time cell analysis (RTCA) neutralization assay**

To determine the neutralizing activity of serum/plasma against SARS and SARS-2, we used real-time cell analysis (RTCA) assay on an xCELLigence RTCA MP Analyzer (ACEA Biosciences Inc.) that measures virus-induced cytopathic effect (CPE) (*3, 4*). Briefly, 50 µL of cell culture medium (DMEM supplemented with 2% FBS) was added to each well of a 96-well E-plate using a ViaFlo384 liquid handler (Integra Biosciences) to obtain background reading. A suspension of 18,000 Vero-E6 cells in 50 µL of cell culture medium was seeded in each well, and the plate was placed on the analyzer. Measurements were taken automatically every 15 min, and the sensograms were visualized using RTCA software version 2.1.0 (ACEA Biosciences Inc). Replication-competent VSV expressing SARS (VSV-SARS-CoV) or SARS-2 spike proteins (VSV-SARS-CoV-2) at 0.01 MOI (~120 PFU per well) was mixed 1:1 with a dilution of serum/plasma or mAb in a total volume of 100 µL using DMEM supplemented with 2% FBS as a diluent and incubated for 1 h at 37°C in 5% $CO_2$. At 16 h after seeding the cells, the virus-mAb mixtures were added in replicates to the cells in 96-well E-plates. Triplicate wells containing virus only (maximal CPE in the absence of mAb) and wells containing only Vero cells in medium (no-CPE wells) were included as controls. Plates were measured continuously (every 15 min) for 48 h to assess virus neutralization. Normalized cellular index (CI) values at the endpoint (48 h after incubation with the virus) were determined using the RTCA software version 2.1.0 (ACEA Biosciences Inc.). Results are expressed as percent neutralization in a presence of respective mAb relative to control wells with no CPE minus CI values from control wells with maximum CPE. RTCA $IC_{50}$ values were determined by nonlinear regression analysis using Prism software.

**Serum IgG isolation and Fab digestion**

For IgG isolation, 1 mL of human serum diluted to 5 mL with PBS was incubated with 500 µL of washed protein G resin (GE Healthcare) overnight at 4°C. The resin was washed three times with PBS and eluted with 10 mL of 0.1 M glycine buffer at pH 2.5. The elute was immediately neutralized with 4 mL of 1M Tris-HCl, pH 8.0 and buffer exchanged to PBS using 100 kDa cutoff Amicon ultrafiltration units. For Fab preparation, 4 mg of concentrated polyclonal IgG samples were incubated with papain-agarose resin (Thermo Fisher Scientific) in digestion buffer (20 mM sodium phosphate, 10 mM EDTA, 20 mM cysteine, pH 7.4) for around 22 h in a 37°C incubator. The digest was removed from the beads and buffer exchanged to PBS. The undigested IgGs were removed by SEC using a Superose 6 increase 10/300 column (GE Healthcare Biosciences). The purified Fabs were concentrated and assessed by Sodium Dodecyl Sulfate - PolyAcrylamide Gel Electrophoresis (SDS-PAGE) for purity.

**Preparation of Fab-spike complexes for ns-EMPEM**

Fab-spike complexes were generated by incubating 20 µg of spike protein with 1 to 1.5 mg of purified polyclonal Fabs overnight at RT. For complexes with the SARS-2 spike, 20 µg of a mixture of spikes (GSAS-2P, GSAS-2P mut2, GSAS-2P mut4 and GSAS-2P mut7) were complexed with 5 mg of polyclonal Fabs overnight at RT. The complexes were purified on a Superose 6 increase 10/300 column using UV absorbance at 215 nm on Akta Pure system (GE Healthcare) running in TBS buffer. The fractions containing spike-Fab complexes were concentrated using 10 kDa cutoff Amicon ultrafiltration units and immediately used for making EM grids.

**Ns-EMPEM sample preparation and data collection**

Spike-polyclonal Fab complexes diluted to approximately 20 µg/mL with TBS were directly deposited onto carbon-coated 400-mesh copper grids (made in house) and stained with 2 % (w/v) uranyl-formate for 90 seconds immediately following sample application. Grids were imaged at 120 keV on Tecnai T12 Spirit with either a 4kx4k Tietz TemCam-F416 detector or with a 4kx4k Eagle CCD (52,000 × magnification at ~1.5 µm under focus). Micrographs were collected using Leginon and the images were transferred to Appion for processing (*5, 6*). Particle stacks were generated in Appion with particles picked using a Difference-of-Gaussians picker (DoG-picker) (*7*). Particle stacks were then transferred to Relion for 2D classification followed by 3D classification to sort classes based on different Fab specificities (*8*). Classes with similar specificities were iteratively assembled and reclassified to generate final reconstructions. A subset of 3D classes with good Fab reconstructions were auto-refined on Relion and used for making composite figures using UCSF Chimera or ChimeraX (*9, 10*). Among the OC43- and

HKU1-reactive antibodies that were detected by EMPEM 2D classes, certain specificities did not refine into 3D reconstructions as a consequence of either low particle numbers for the Fab class, orientation bias on the grid, or due to polyclonal Fab specificities targeting the same epitope. 3D refined maps were successfully generated for all OC43-reactive donors and 1of 2 HKU1-reactive donors.

**Cryo-EMPEM sample preparation**

For cryo-EMPEM studies with OC43 spike, 50 µg of the spike protein was complexed with 3 mg of purified polyclonal fab from each donor. The complex was incubated overnight at RT and purified as described above. For OC43 spike-polyclonal Fab complexes made with donor 269 and donor 1051, 3.5 µl of complex at 0.5 µg/mL and 0.7 µg/mL were mixed with 0.5 µl of 0.04 mM Lauryl maltose neopentyl glycol (LMNG) solution immediately before sample deposition onto 1.2/1.3 300-mesh UltraAuFoil grids (EMS). For OC43 spike-polyclonal Fab complex from donor 1412, 3.5 µl of spike-Fab complex at 0.5 µg/mL was mixed with 0.5 µl of 0.48 mM n-dodecyl-B-D-maltopyranoside (DDM) solution before sample deposition onto 1.2/1.3 300-mesh UltraAuFoil grids (EMS). Grids were plasma cleaned for 7 seconds prior to sample deposition using Gatan Solarus 950 Plasma system ($Ar/O_2$ gas mixture). Following sample application, grids were blotted for 4 seconds before being plunged into liquid nitrogen-cooled liquid ethane using a Vitrobot mark IV (Thermo Fisher Scientific).

For cryo-EMPEM studies with SARS-2 spike, 40 µg of spike mixture comprising of equal ratios of HP-GSAS, HP-GSAS Mut2, HP-GSAS Mut4 and HP-GSAS Mut7 spikes were incubated overnight at RT with 10 mg of purified polyclonal Fabs (5 mg each from donor 1988 and 1989) before purifying the complexes. The complexes were mixed with LMNG immediately prior to sample deposition onto plasma cleaned Quantifoil 1.2/1.3 grids (EMS) that were blotted for 3 seconds and plunge-frozen in liquid ethane using a Vitrobot.

**Cryo-EM data collection and processing**

Micrographs were collected through Leginon software on a FEI Titan Krios operating at 300 keV mounted with a Gatan K2 direct-electron detector. The collection parameters are described in table S1. MotionCor2 was used for alignment and dose weighting of the frames and micrographs were transferred to CryoSPARC 2.9 for initial processing (*11*, *12*). CTF estimations were performed using GCTF and micrographs selected using the Curate Exposures tool in CryoSPARC based on CTF resolution estimates (cutoff 5 Å) for downstream particle picking, extraction and iterative rounds of 2D classification and selection of intact spike trimers (*13*). The clean particle stacks were then transferred to Relion for 3D refinement and different antibody classes were sorted using the focused classification protocol described previously (*14*). The larger datasets obtained for donors 269 (~2.2 million particles post symmetry expansion) and 1051 (~2.1 million particles post symmetry expansion) resulted in reconstructions ranging from 3-3.5 Å resolution while the smaller dataset (~0.7 million particles post symmetry expansion) collected for 1412 resulted in reconstructions between 4-8 Å resolution. For SARS- 2 cryoEMPEM, 5 Fab-spike complexes were reconstructed at resolutions between 3.9-4.4 Å from a dataset of ~1.1 million particles post symmetry expansion. The data collection parameters and the processing workflow are summarized in Table S1 and Figures S2, S3, S4 and S8.

**Model building and refinement.** Initial model building into OC43 spike-antibody 3D maps was performed manually in Coot using PDB 6OHW as a template for the spike. Fabs are represented as poly-alanine backbone models, as their exact sequence is unknown. Iterative rounds of Rosetta relaxed refinement and manual Coot refinement were applied to generate the final models (*15-17*). EMRinger and MolProbity metrics were calculated following each round of Rosetta refinement to evaluate and identify the best refined models (*18, 19*). Phenix comprehensive validation was performed on the final models. To prepare sapienic acid for modeling, PDB and CIF ligand definition files were created using Phenix eLBOW by providing the SMILES string for PubChem CID: 5312419 (Sapienic acid) (*20*). The coordinates were manually placed into their map densities in the spike and refined using Coot.  Final map and model statistics are summarized in Tables S2 and S4

**Mass-spectrometry.** Mass spectrometry was performed as previously described to confirm the identity of sapienic acid in the apo-OC43 spike protein (*21*). Briefly, Acetonitrile was used to precipitate the spike protein and extract the fatty acid followed by ESI-TOF high accuracy mass spectrometry to screen for the compound of interest in the 250 to 300 m/z range. We obtained a single hit at molecular weight of 254 g/mol that matches with sapienic acid.

#### Figures

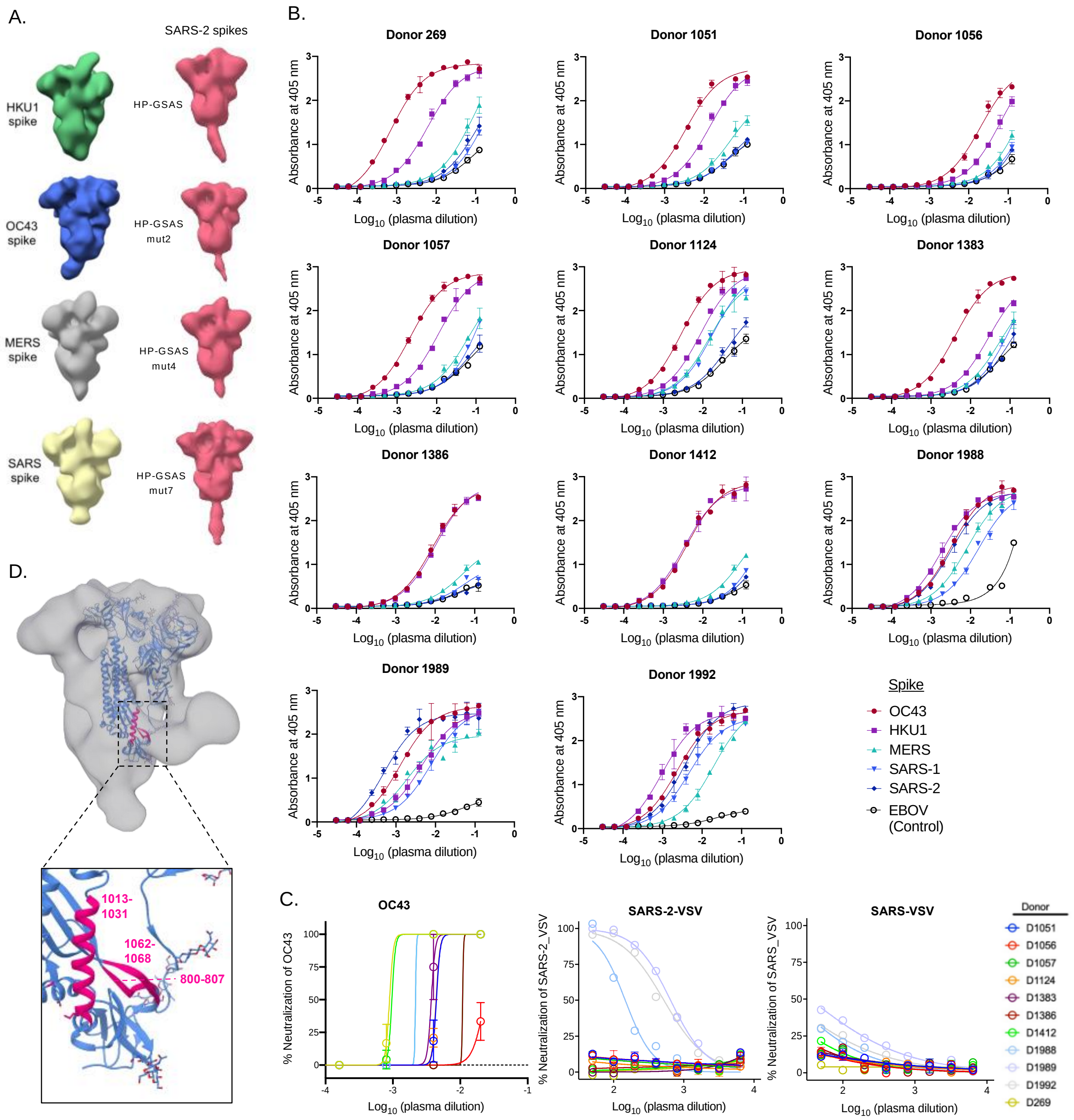


**Figure S1. Serum/plasma antibody responses to β-CoV spikes (A)** ns-EM reconstructions of pre-fusion stabilized spike ectodomains from OC43 (green), HKU1 (blue), MERS (grey), SARS (yellow) and SARS-2 (pink). For SARS-2, a total of four constructs are shown: a base construct (HP-GSAS) with 6 stabilizing proline substitutions and furin cleavage site modification and 3 other constructs with additional disulphide linkages, HP-GSAS Mut2 (S383C and D985C), HP-GSAS Mut4 (A570C and L966C) and HP-GSAS Mut7 (V705C and T883C). **(B)** Binding curves for 11 donor plasma/serum samples against β-CoV spikes as determined by ELISA. Ebolavirus glycoprotein (EBOV GP) was used as a control to determine baseline non-specific serum binding **(C)** Serum or plasma neutralization curves for 11 donors against OC43 virus or VSV pseudotyped SARS or SARS-2 viruses. **(D)** ns-EM 3D reconstruction (grey) of OC43 spike in complex with a S2 antibody from donor 1051. One spike protomer (blue) in the model (PDB: 6OHW) is shown docked into the EM density and a zoom-in of the antibody footprint in pink is shown in the bottom panel (*15*).

A. Cryo-EM analysis of HCoV-OC43 spike + polyclonal Fab from donor 269

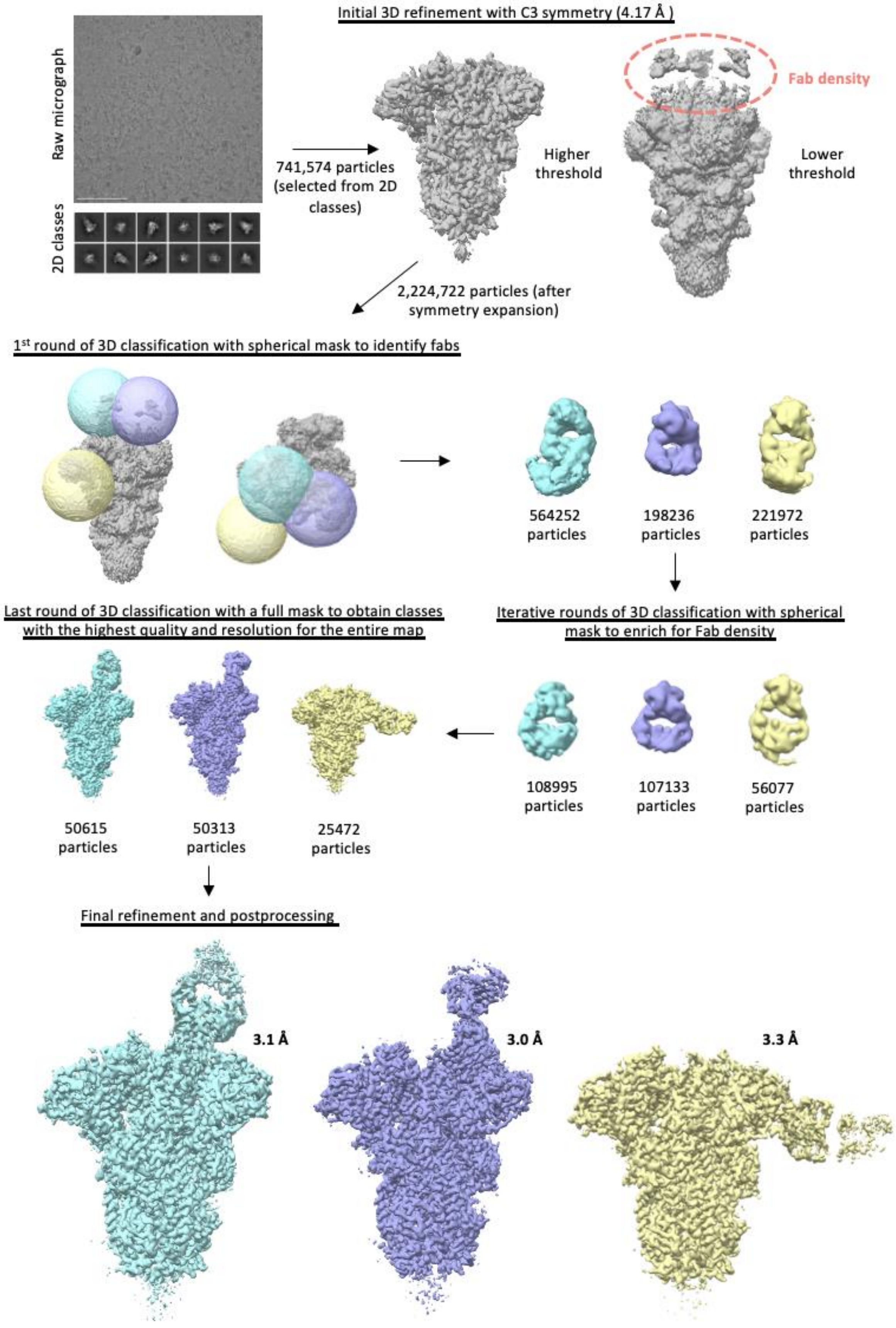

**Figure S2. Schematic representation of the cryo-EMPEM processing workflow for OC43 spike complexed with polyclonal Fabs from donor 269**. The focused classification approach used for generating Fab-spike reconstructions is shown in steps.

A. Cryo-EM analysis of HCoV-OC43 spike + polyclonal Fab from donor 1051

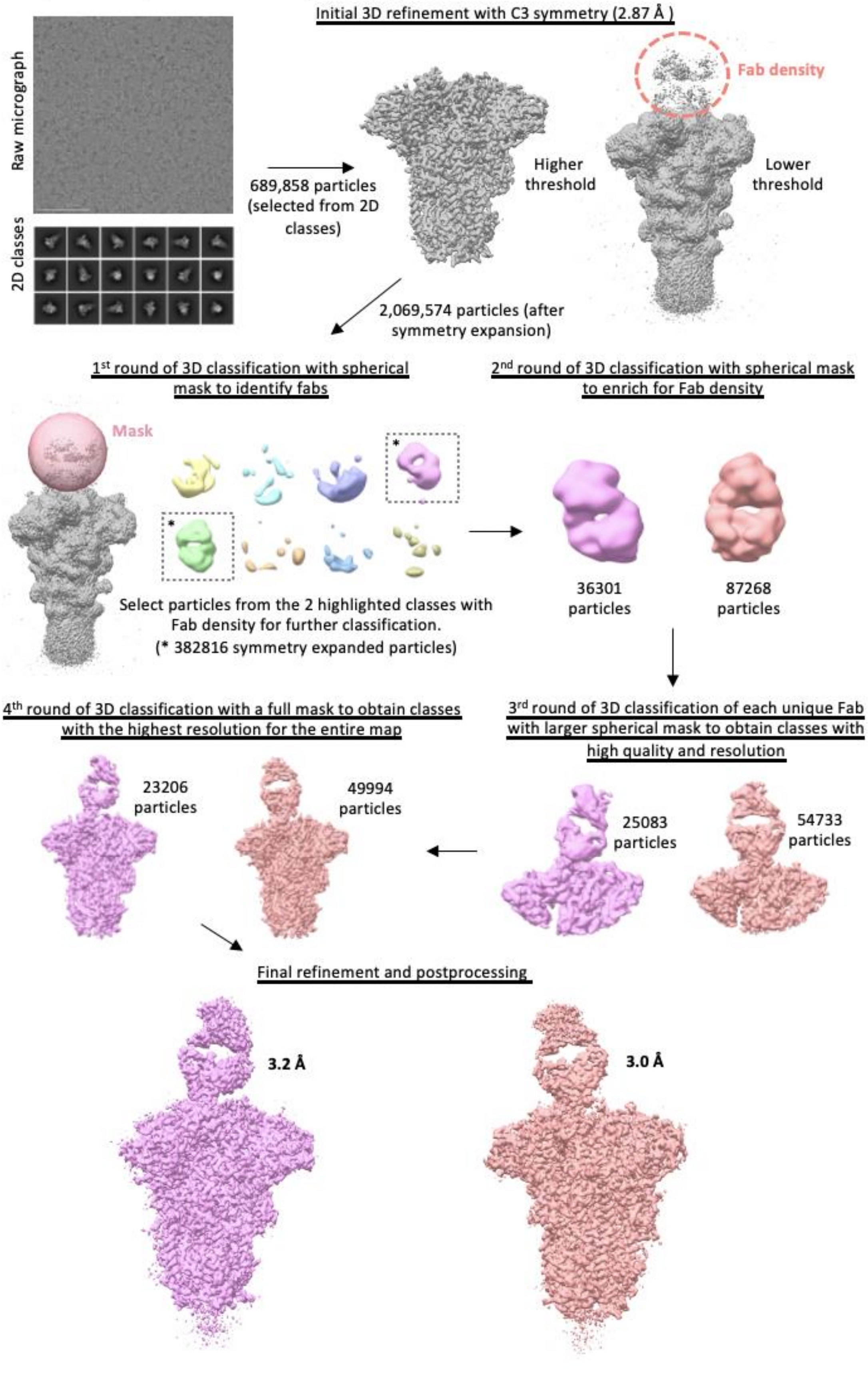

**Figure S3. Schematic representation of the cryo-EMPEM processing workflow for OC43 spike complexed with polyclonal Fabs from donor 1051**. The focused classification approach used for generating Fab-spike reconstructions is shown in steps.

**A. Cryo-EM analysis of HCoV-OC43 spike + polyclonal Fab from donor 1412**

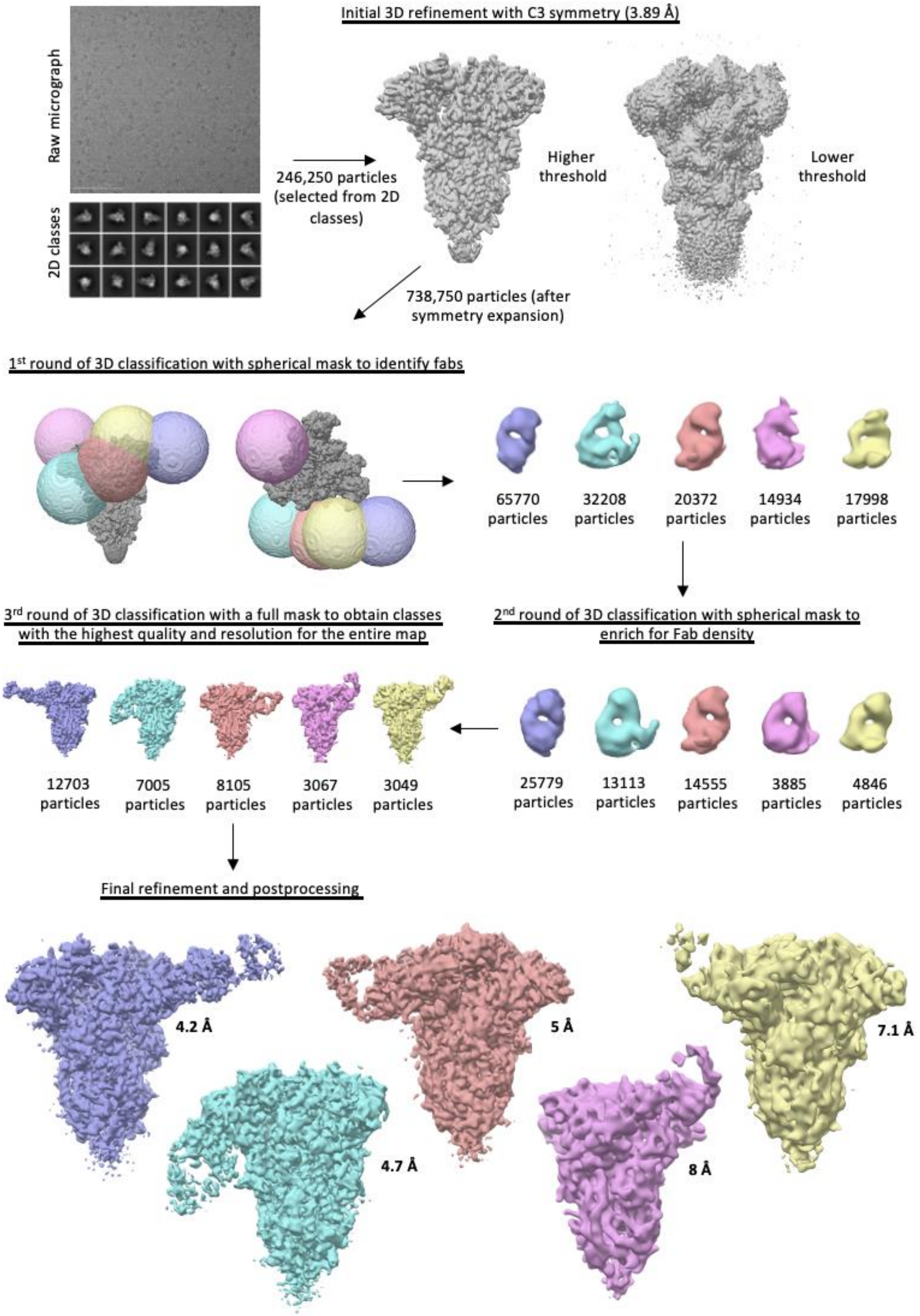

**Figure S4. Schematic representation of the cryo-EMPEM processing workflow for OC43 spike complexed with polyclonal Fabs from donor 1412**. The focused classification approach used for generating Fab-spike reconstructions is shown in steps.

A.

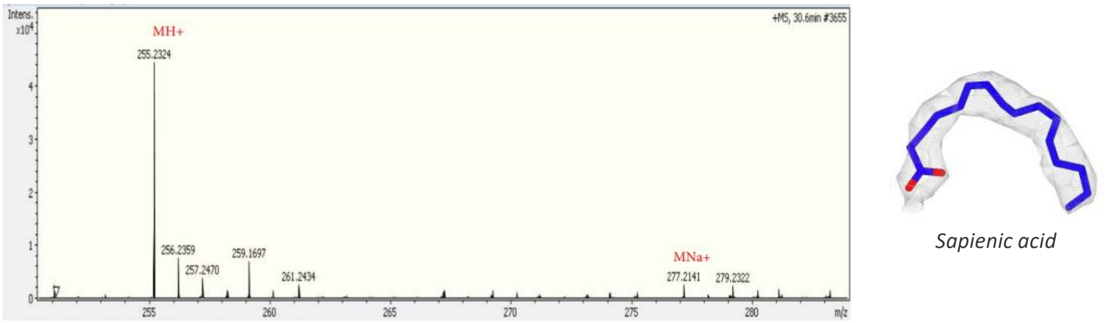

Intens.
x10^4
+MS, 30.6min #3655
MH+
255.2324
256.2359
257.2470
259.1697
261.2434
MNa+
277.2141
279.2322
m/z
Sapienic acid


B.

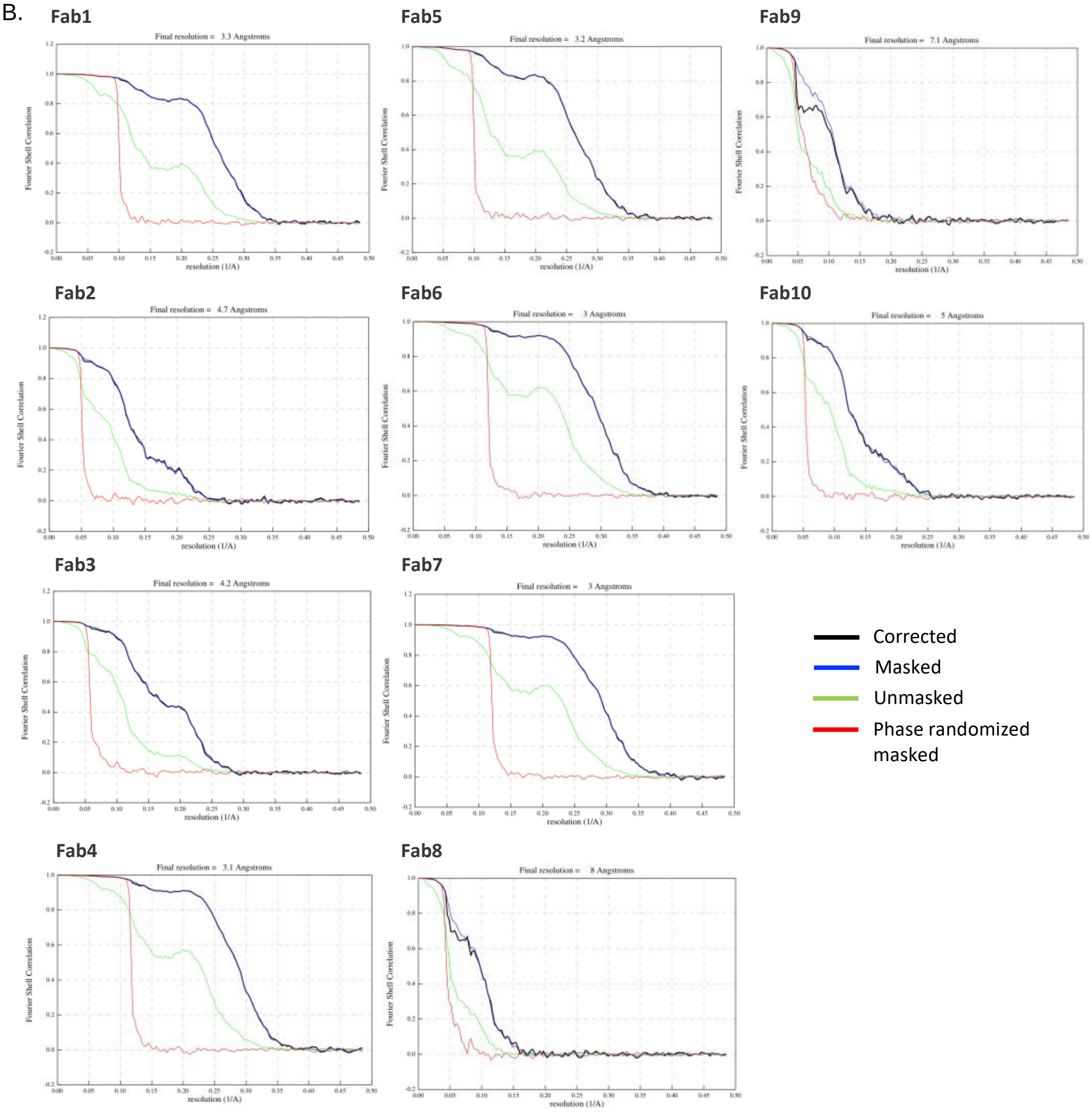

Fab1
Final resolution = 3.3 Angstroms
Fab5
Final resolution = 3.2 Angstroms
Fab9
Final resolution = 7.1 Angstroms
Fab2
Final resolution = 4.7 Angstroms
Fab6
Final resolution = 3 Angstroms
Fab10
Final resolution = 5 Angstroms
Fab3
Final resolution = 4.2 Angstroms
Fab7
Final resolution = 3 Angstroms
Fab4
Final resolution = 3.1 Angstroms
Fab8
Final resolution = 8 Angstroms
Fourier Shell Correlation
resolution (1/A)
Corrected
Masked
Unmasked
Phase randomized masked

**Figure S5. Cryo-EMPEM structure analysis and validation (A)** Left panel shows mass spectrometry results for HCoV-OC43 spike showing a species with MW of 255 g/mol which corresponds to the protonated form of sapienic acid (MW = 254 g/mol). The right panel shows sapienic acid colored in blue surrounded by its corresponding cryo-EM map density shown as a grey mesh. Sapienic acid has a kink at position 6, originating from a cis double bond, that perfectly recapitulates the density in our HCoV-OC43 cryo-EM maps. **(B)** FSC curves for HCoV-OC43 spike-Fab cryo-EMPEM reconstructions with C1 symmetry imposed.

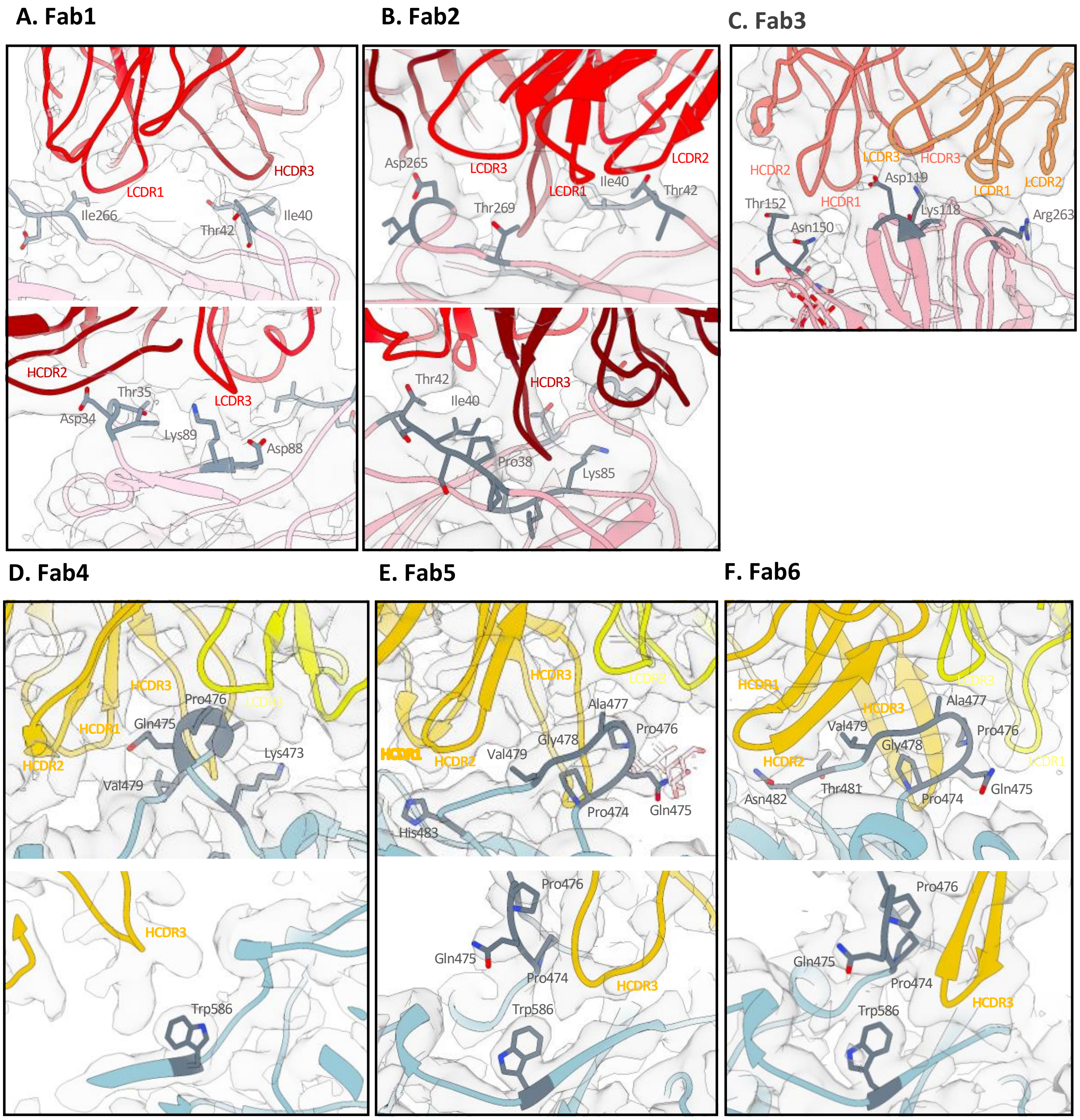


**Figure S6. Epitope-paratope interactions for polyclonal Fabs 1-6 targeting the OC43 spike NTD or CTD.** Ribbon representation of atomic models of OC43 spike-Fab complexes docked into their respective cryo-EM maps with close-up views of epitope-paratope interactions for (A) Fab1, (B) Fab2, (C) Fab3, (D) Fab4, (E) Fab5 and (F) Fab6. The heavy or light chains are colored dark red or light red for NTD-site 1 Fabs (Panels A and B), colored coral or orange for the NTD-site 2

Fab (Panel C), colored mustard or yellow for CTD Fabs (Panel D, E and F) and the spike is colored pink (Panels A, B and C) or blue (Panels D, E and F). The interacting residues are colored grey.

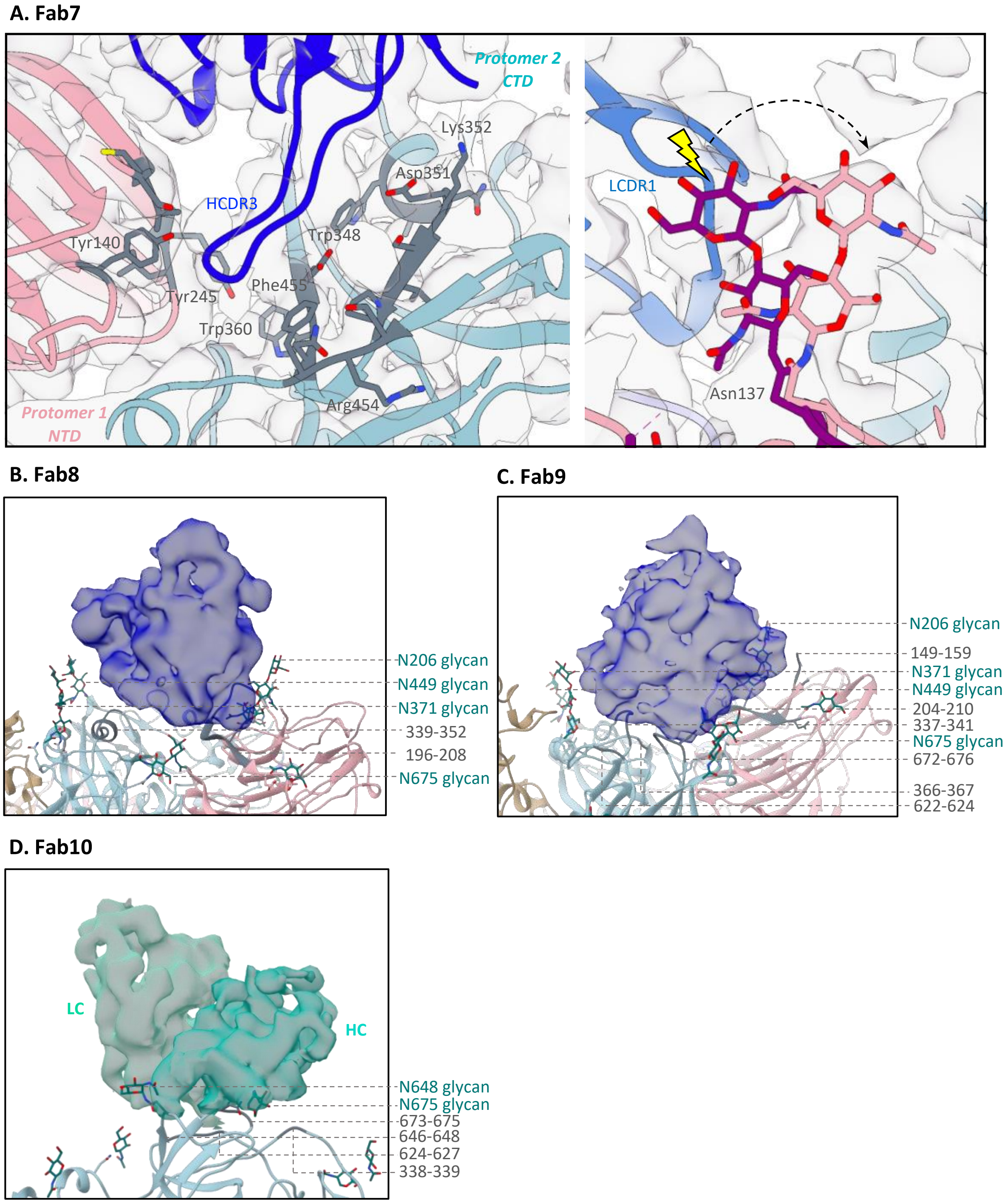

**Figure S7. Epitope-paratope interactions for polyclonal Fabs 7-10 targeting the OC43 spike inter-protomeric interfaces or SD1. (A)** Ribbon representation of atomic model for OC43 spike-Fab7 complex docked into the cryo-EM map with close-up views of epitope-paratope interactions. The Fab heavy and light chains are colored dark blue and light blue, respectively. The adjacent spike protomers are colored pink and blue with the interacting residues colored grey. The right panel shows displacement of N137 glycan (light pink) by LCDR3 compared to the published structure shown in magenta (PDB 6OHW)(*15*). **(B and C)** Close-up views of epitope-paratope interactions for OC43 spike- **(B)** Fab8 and **(C)** Fab9 complexes displaying the cryo-EM density corresponding to the Fab in blue with ribbon representation of spike protomers in blue, pink or tan and interacting residues in grey. **(D)** Close-up view of epitope-paratope interaction for OC43 spike-Fab10 complex displaying the cryo-EM density corresponding to Fab heavy chain and light chain in shades of cyan and ribbon representation of a spike protomer in blue with interacting residues in grey. Glycans are colored teal in panels B, C and D.

**A. Cryo-EM analysis of SARS-2 spikes + pooled polyclonal Fabs from donor 1988 and 1989**

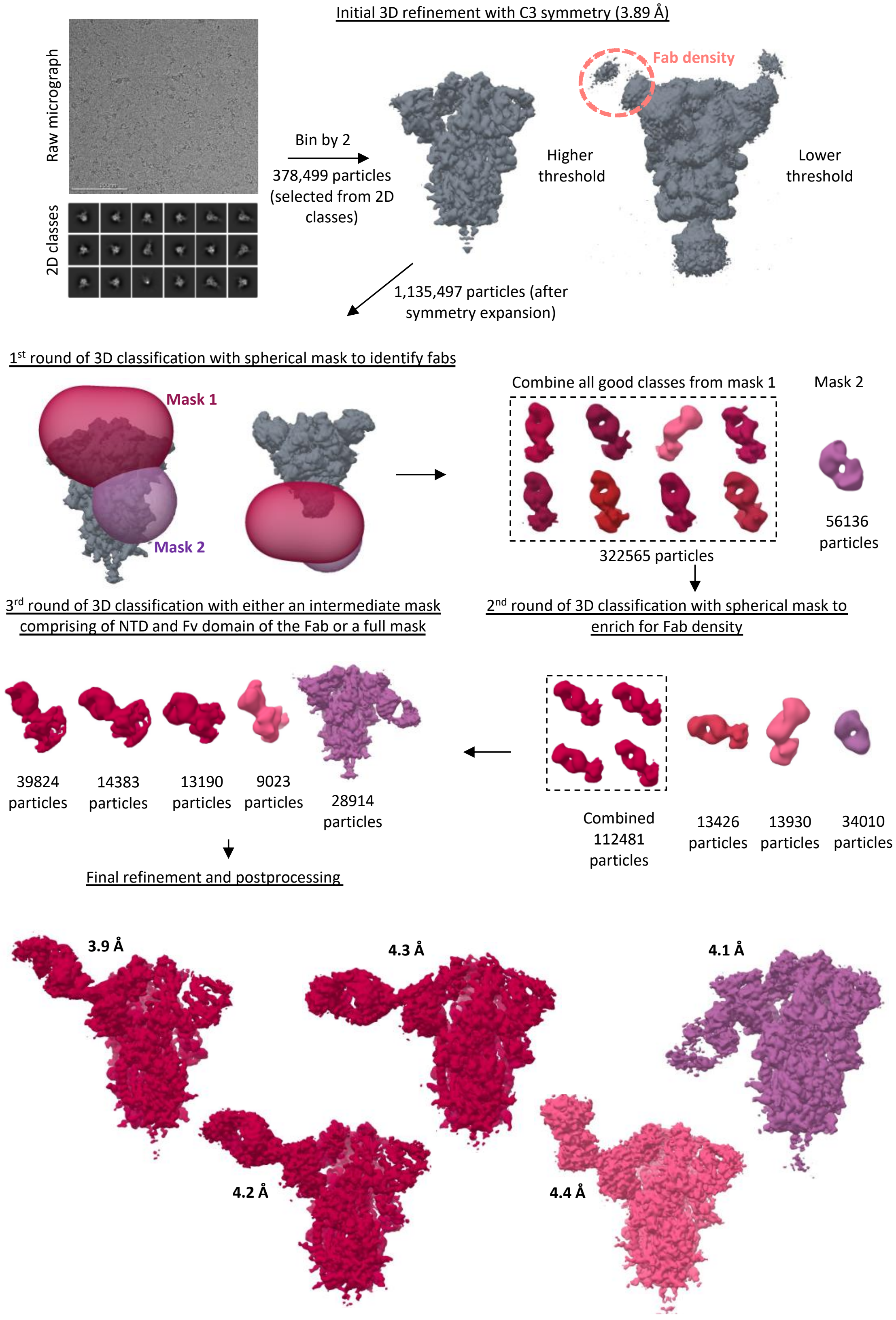

**Figure S8. Schematic representation of the cryo-EMPEM processing workflow for SARS-2 spikes complexed with polyclonal Fabs from donors 1988 and 1989**. The focused classification approach used for generating Fab-spike reconstructions is shown in steps.

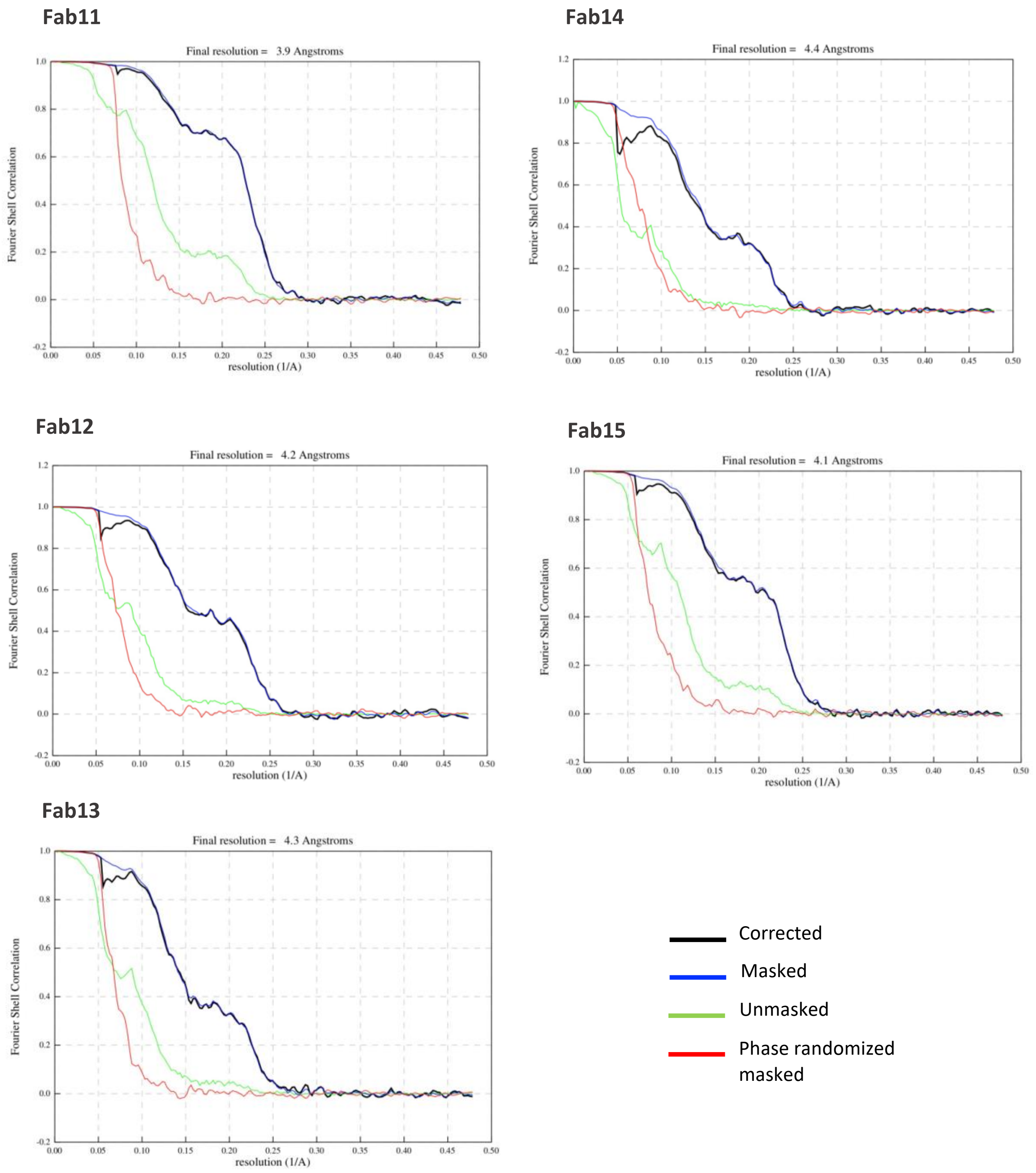


**Figure S9. FSC curves for SARS-2 spike-Fab cryo-EMPEM reconstructions with C1 symmetry imposed.**

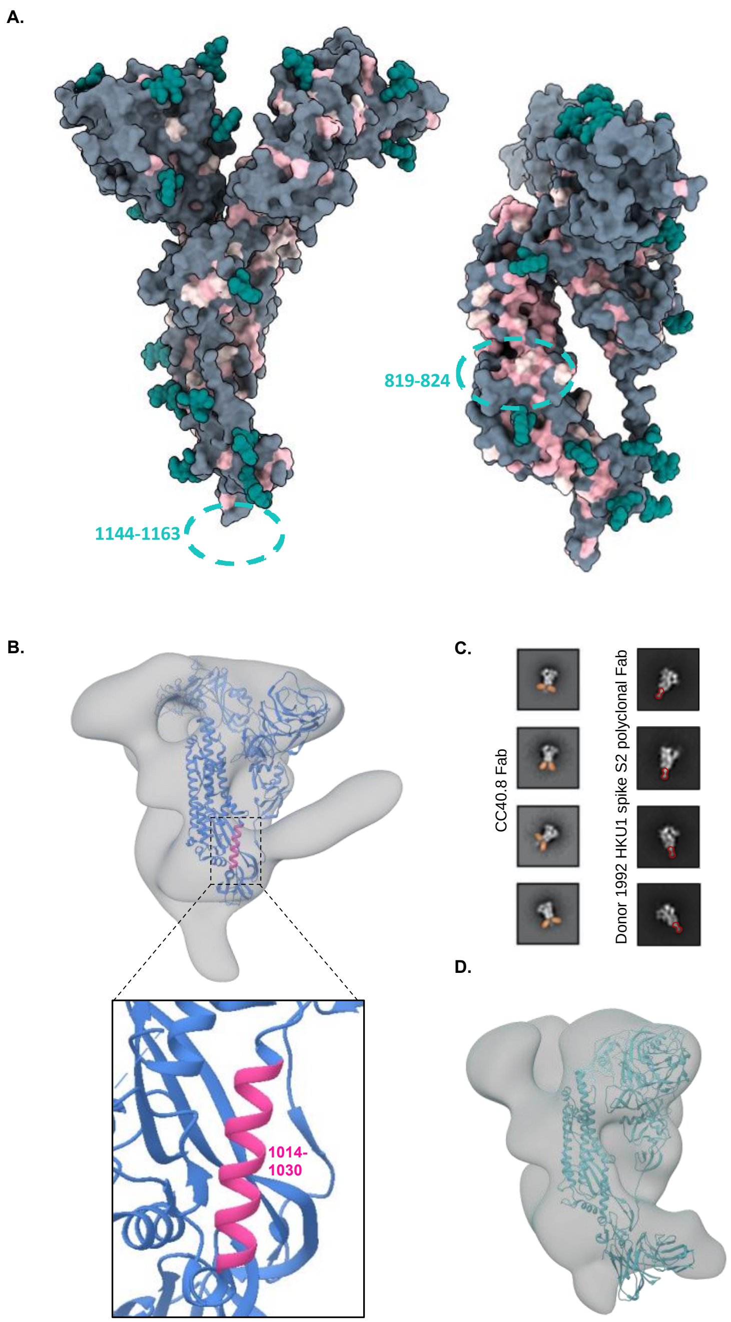


**Figure S10. β-CoV cross-reactive responses in SARS-2 convalescent donors. (A)** Surface representation of SARS-2 spike monomer (PDB 7JJI) colored grey and glycans colored teal. Residues on SARS-2 spike that are either identical to or have a conserved substitution in at least 3

of the 4 other β-CoVs, OC43, HKU1, SARS and MERS are colored in pink and light pink, respectively. Conserved residues of interest from published studies (*22*, *23*) are depicted on the model for reference. **(B)** ns-EM 3D reconstruction (grey) of OC43 spike in complex with a S2 antibody from donor 1989. One spike protomer (blue) in the model (PDB: 6OHW)(*15*) is shown docked into the EM density and a zoom-in of the antibody footprint in pink is shown in the bottom panel. **(C)** ns-EM 2D class averages for HKU1 spike complexed with either Fab CC40.8 (*24*) or a S2 antibody from donor 1992; the Fabs are false-colored orange and red, respectively. (D) Nn-EM 3D reconstruction (grey) of MERS spike in complex with a S2 antibody from donor 1989. One MERS spike protomer bound to G4 Fab colored in green (PDB: 5W9M)(*25*) is shown docked into the EM density for comparison.

#### Table S1. Cryo-EM data collection

| Data collection | HCoV-OC43-269 | HCoV-OC43-1051 | HCoV-OC43-1412 | SARS-CoV-2-1988/1989 |
|---|---|---|---|---|
| Microscope | FEI Titan Krios | FEI Titan Krios | FEI Titan Krios | FEI Titan Krios |
| Voltage (kV) | 300 | 300 | 300 | 300 |
| Detector | Gatan K2 Summit | Gatan K2 Summit | Gatan K2 Summit | Gatan K2 Summit |
| Recording mode | Counting | Counting | Counting | Counting |
| Nominal magnification | 29,000 | 29,000 | 29,000 | 29,000 |
| Movie micrograph pixel size (Å) | 1.03 | 1.03 | 1.03 | 1.045 |
| Dose rate ($e^-$/[(camera pixel)*s]) | 5.6 | 5.7 | 5.3 | 7.2 |
| Number of frames per movie micrograph | 38 | 37 | 40 | 30 |
| Frame exposure time (ms) | 250 | 250 | 250 | 250 |
| Movie micrograph exposure time (s) | 9.5 | 9.25 | 10 | 7.5 |
| Total dose ($e^-$/Å$^2$) | 50 | 50 | 50 | 50 |
| Defocus range (µm) | -0.8 to -1.5 | -0.8 to -1.8 | -0.3 to -1.3 | -0.7 to -1.7 |
| Number of movie micrographs | 4370 | 2575 | 2321 | 5405 |

### Table S2. Refinement and model building statistics for HCoV-OC43-Fab complexes

| Map | HCoV-OC43-Fab1 | HCoV-OC43-Fab2 | HCoV-OC43-Fab3 | HCoV-OC43-Fab4 | HCoV-OC43-Fab5 | HCoV-OC43-Fab6 | HCoV-OC43-Fab7 | HCoV-OC43-Fab8 | HCoV-OC43-Fab9 | HCoV-OC43-Fab10 |
|---|---|---|---|---|---|---|---|---|---|---|
| EMDB | EMD-24968 | EMD-24969 | EMD-24970 | EMD-24989 | EMD-24990 | EMD-24991 | EMD-24992 | EMD-24993 | EMD-24994 | EMD-24995 |
| Donor | 269 | 1412 | 1412 | 269 | 1051 | 1051 | 269 | 1412 | 1412 | 1412 |
| Number of molecular projection images in map | 25,472 | 7,005 | 12,703 | 50,615 | 25,083 | 54,733 | 50,313 | 3,885 | 4,846 | 14,555 |
| Symmetry | C1 | C1 | C1 | C1 | C1 | C1 | C1 | C1 | C1 | C1 |
| Map resolution (FSC 0.143; Å) | 3.3 | 4.7 | 4.2 | 3.1 | 3.2 | 3.0 | 3.0 | 8.0 | 7.1 | 5.0 |
| Map sharpening B-factor (Å$^2$) | -64.7 | -137.8 | -104.5 | -76.9 | -63.4 | -70.3 | -67.9 | -415.6 | -242.8 | -105.1 |
| **Structure Building and Validation** | | | | | | | | | | |
| *Number of residues in deposited model* | | | | | | | | | | |
| Amino acids | 3756 | 3739 | 3755 | 3740 | 3790 | 3769 | 3738 | | | |
| Carbohydrates | 56 | 55 | 55 | 56 | 85 | 62 | 58 | | | |
| Sapienic acid | 3 | 3 | 3 | 3 | 3 | 3 | 3 | | | |
| MolProbity score | 1.02 | 1.07 | 1.03 | 0.80 | 0.76 | 0.76 | 0.69 | | | |
| Clashscore | 1.85 | 1.93 | 1.62 | 0.54 | 0.78 | 0.84 | 0.50 | | | |
| EMRinger score | 3.62 | 0.83 | 2.06 | 4.5 | 4.32 | 4.9 | 4.74 | | | |
| *RMSD from ideal* | | | | | | | | | | |
| Bond length (Å) | 0.018 | 0.019 | 0.020 | 0.022 | 0.022 | 0.023 | 0.021 | | | |
| Bond angles (˚) | 1.674 | 1.718 | 1.704 | 1.708 | 1.654 | 1.645 | 1.749 | | | |
| *Ramachandran plot* | | | | | | | | | | |
| Favored (%) | 97.7 | 97.4 | 97.4 | 97.4 | 97.9 | 98 | 97.9 | | | |
| Allowed (%) | 2.3 | 2.6 | 2.6 | 2.6 | 2.1 | 2 | 2.1 | | | |
| Outliers (%) | 0 | 0 | 0 | 0 | 0 | 0 | 0 | | | |
| Side chain rotamer outliers (%) | 0.36 | 0 | 0 | 0.23 | 0.42 | 0.26 | 0.16 | | | |
| PDB | 7SB3 | 7SB4 | 7SB5 | 7SBV | 7SBW | 7SBX | 7SBY | | | |

#### Table S3. Predicted CDR lengths for HCoV-OC43 Fabs

| Fab | Heavy Chain | | | Light Chain | | |
|---|---|---|---|---|---|---|
| | HCDR1 | HCDR2 | HCDR3 | LCDR1 | LCDR2 | LCDR3 |
| Fab1 | 8 | 11 | 14 | 8 | 3 | 9 |
| Fab2 | 8 | 11 | 14 | 6 | 2 | 9 |
| Fab3 | 8 | 9 | 10 | 6 | 3 | 8 |
| Fab4 | 10 | 11 | 15 | 4 | 3 | 9 |
| Fab5 | 8 | 11 | 18 | 6 | 2 | 9 |
| Fab6 | 8 | 10 | 19 | 12 | 3 | 9 |
| Fab7 | 8 | 13 | 15 | 7 | 3 | 5 |

#### Table S4. Refinement and model building statistics for SARS-2-Fab complexes

| Map | SARS-CoV-2-Fab11 | SARS-CoV-2-Fab12 | SARS-CoV-2-Fab13 | SARS-CoV-2-Fab14 | SARS-CoV-2-Fab15 |
|---|---|---|---|---|---|
| EMDB | EMD-24996 | EMD-24997 | EMD-24998 | EMD-24999 | EMD-25000 |
| Donors | 1988/1989 | 1988/1989 | 1988/1989 | 1988/1989 | 1988/1989 |
| Number of molecular projection images in map | 39,824 | 14,383 | 13,190 | 9,023 | 28,914 |
| Symmetry | C1 | C1 | C1 | C1 | C1 |
| Map resolution (FSC 0.143; Å) | 3.9 | 4.2 | 4.3 | 4.4 | 4.1 |
| Map sharpening B-factor (Å$^2$) | -76.2 | -93.1 | -81.2 | -90.0 | -79.3 |
